# Impaired lysosomal acidification triggers iron deficiency, necrotic cell death and inflammation *in vivo*

**DOI:** 10.1101/710798

**Authors:** King Faisal Yambire, Christine Rostosky, Takashi Watanabe, David Pacheu-Grau, Sylvia Torres-Odio, Angela Sanchez-Guerrero, Ola Senderovich, Esther G. Meyron-Holtz, Ira Milosevic, Jens Frahm, Phillip West, Nuno Raimundo

## Abstract

Lysosomal acidification is a key feature of healthy cells. Inability to maintain lysosomal acidic pH is associated with aging and neurodegenerative diseases. However, the mechanisms elicited by impaired lysosomal acidification remain unknown. We show here that inhibition of lysosomal acidification triggers cellular iron deficiency, which results in impaired mitochondrial function and necrotic cell death. These effects are recovered by supplying iron via a lysosome-independent pathway. Notably, iron deficiency is sufficient to trigger inflammatory signaling in cultured primary neurons. Using a mouse model of impaired lysosomal acidification, we observed a robust iron deficiency response in the brain, verified by *in vivo* magnetic resonance imaging. Furthermore, the brains of these mice present a pervasive inflammatory signature associated with instability of mitochondrial DNA (mtDNA), both corrected by supplementation of the mice diet with iron. Our results highlight a novel mechanism linking lysosomal dysfunction, mitochondrial malfunction and inflammation *in vivo*.

## INTRODUCTION

Lysosomal function is now recognized as a key factor in cellular and tissue health. Recessive mutations in genes encoding lysosomal proteins result in over 50 severe lysosomal storage diseases, and carriers of these mutations are at risk of many neurodegenerative diseases, such as Parkinson’s (Straud et al., 2010), Alzheimer’s (Garcia-Rodriguez et al., 2015), amyotrophic lateral sclerosis (Farsi et al., 2018), frontotemporal lobar degeneration (Lie and Nixon, 2019), among others (Bowman et al., 1988; Xie et al., 2004). The lysosomes are now recognized as key players in cellular signaling and nutrient sensing, in addition to their roles as terminal platform of autophagy and endocytosis (Perera and Zoncu, 2016). Lysosomes are also involved in the intracellular partitioning of several cellular building blocks, such as amino acids, cholesterol and sphingomyelin, as well as metals including calcium (Ca) and iron (Fe) (Lim and Zoncu, 2016).

There are different populations of lysosomes in each cell, which can be distinguished by their position, size, acidification and reformation properties (Durrbaum et al., 2014). Yet, most lysosomal functions rely on the acidification of the lysosomal lumen, as the vast majority of the enzymes residing in the lysosome have an optimal function in the pH range of 4.5-5 (Perera and Zoncu, 2016). The acidic pH of the lysosomal lumen is the result of an electrochemical gradient maintained mostly by the vacuolar ATPase (v-ATPase) (Wang and Semenza, 1993), with contribution from the chloride channel CLC-7 (Hamdi et al., 2016; Santaguida et al., 2015). The v-ATPase is a multisubunit complex with two domains, the membrane-associated V_O_ and the soluble hydrolytic V_1_, which hydrolyzes ATP to pump protons to the lysosomal lumen against a concentration gradient (Rouault, 2015). The existence of specific v-ATPase inhibitors such as bafilomycin A1 (interacts with the V_o_ ring, inhibiting proton translocation) and saliphenylhalamide (saliphe; locks v-ATPase in an assembled state) provides pharmacological tools to assess loss of v-ATPase activity (Bowman et al., 1988; Garcia-Rodriguez et al., 2015; Xie et al., 2004).The v-ATPase exists also in other cellular organelles, such as endosomes, Golgi complex and secretory vesicles (Farsi et al., 2018), but the effects of v-ATPase loss-of-function most severely relate to its role in lysosomal acidification (Lie and Nixon, 2019).

Decreased activity of the v-ATPase has been linked to age-related decrease in lysosomal function and neurodegenerative diseases (Nixon, 2013; Wang and Semenza, 1993). It has also been shown that impaired acidification of the yeast vacuole, the evolutionary ancestor of lysosomes, results in decreased mitochondrial function and accelerated aging (Mancias et al., 2014). Nevertheless, the mechanisms by which impaired lysosomal acidification result in aging and disease remain poorly characterized. In addition, the v-ATPase has been recognized as a therapeutic target in cancer, given that its inhibition positively correlates with decreased tumor mass (Allen et al., 2013). Therefore, it is pivotal to characterize the mechanisms underlying the cellular response to v-ATPase inhibition. Importantly, the potent inhibition of v-ATPase causes cell death, but the underlying mechanisms also remain unclear. The understanding of which signaling pathways are elicited in response to v-ATPase inhibition would allow the definition of therapeutic targets, for example for neurodegenerative diseases and for cancer. Here, we show that inhibition of the v-ATPase results in impairment of lysosomal iron metabolism which causes iron deficiency in cytoplasm and mitochondria. This results in activation of the pseudo-hypoxia response, loss of mitochondrial function and necrotic cell death. These effects were all ablated by iron repletion in a form that can be imported across the plasma membrane, independently of the lysosomes. Notably, iron deficiency is sufficient to trigger inflammatory signaling in cultured neurons as well as *in vivo*. A mouse model of impaired lysosomal acidification shows iron deficiency, activation of the pseudo-hypoxia response, and pervasive inflammation, all detectable long before the disease onset. All these phenotypes could be rescued by increasing the levels of iron in the animal diet.

## RESULTS

### v-ATPase inhibition triggers hypoxia-inducible factor-mediated response

With the goal of identifying signaling events caused by impaired organelle acidification, we analyzed several transcriptome datasets of cells treated with the v-ATPase inhibitor bafilomycin. These datasets include bafilomycin treatment of HeLa cells (GSE16870) (Straud et al., 2010), colon carcinoma cells (GSE47836) (Durrbaum et al., 2014) and retinal pigment epithelial cells (GSE60570) (Santaguida et al., 2015). We performed multi-dimensional transcriptome analysis in these datasets, aiming at the identification of signaling pathways, networks and transcription factors (Murdoch et al., 2016; Raimundo et al., 2012; Raimundo et al., 2009; Schroeder et al., 2013; Tyynismaa et al., 2010; West et al., 2015; Yambire et al., 2019). We reasoned that those transcription factors (TF) showing similar behavior in the three datasets of bafilomycin-treated cells would be the main regulators of the response to loss of acidification, independently of the cell type. Therefore, we crossed the TF list associated with each dataset, to determine which of those were involved in all three datasets, and found eight common TF, of which seven were predicted as active and one as repressed (Figure 1A). These TF are associated with autophagy (NUPR1), cholesterol homeostasis (SREBF1, SREBF2), hypoxia response (HIF-1α and EPAS1, which is also known as HIF-2α) and diverse stress responses (p53, myc, FoxO3a), and form a highly interconnected network (Figure 1B). To determine which biological processes were associated with these TF, and identify which of them were most upstream, we performed a pathway analysis using Metascape, and found that the most affected processes dealt with cellular response to hypoxia (Figure 1C).

**Figure 1.**
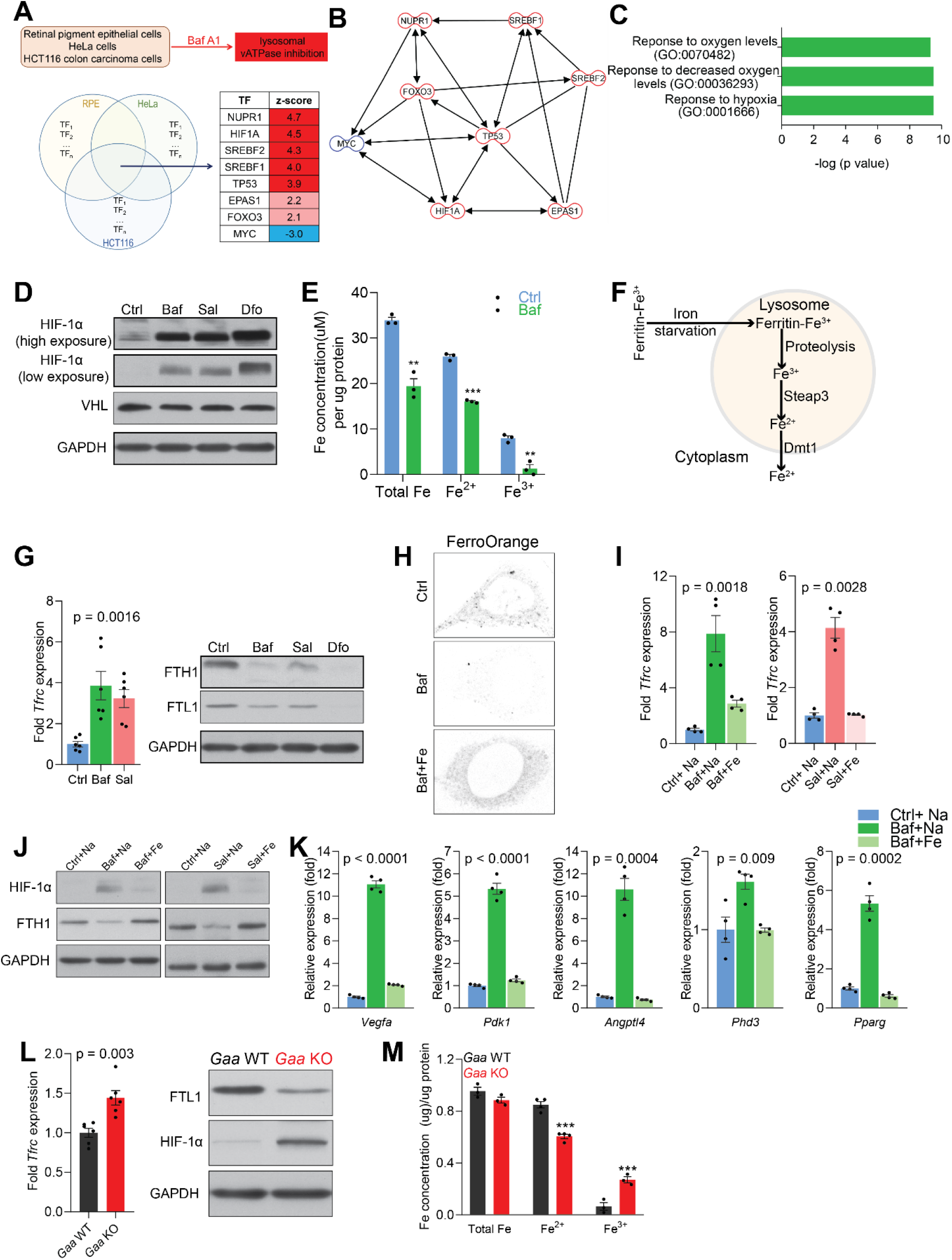
Lysosomal v-ATPase inhibition triggers iron-dependent HIF-1α activation. (A) Venn diagram illustrating common significantly changed upstream regulators (shown as a table) mediating differential gene expression in bafilomycin (Baf)-treated cells in three different transcriptome datasets from HeLa, HCT116 and RPE cells. (B) Transcription factor network of the significantly changed upstream regulators found in Figure 1A. (C) Metascape gene set enrichment analyses show significant overrepresentation of the transcription factors (Figure 1A) in the hypoxia response. (D) Immunoblot of HIF-1α and VHL with GAPDH as loading control in cell lysates of Bafilomycin- and Saliphenylhalamide (Sal)-treated cells. Deferoxamine is used as a positive control for the iron deficiency response. (E) Total, ferrous and ferric iron concentrations in cells treated with or without Bafilomycin for 24 hours. Results are summarized as mean±SEM of three experimental measures shown as black dots. **p < 0.01; ***p <0.001, unpaired two-tailed t-test. (F) Illustration of the intralysosomal pathway of iron homeostasis handling (G) Increased *Tfrc* transcript levels in Baf- and Sal-treated cells relative to untreated cells. Western blot showing decreased FTH1 and FTL1 protein levels in Baf- and Sal-treated cells (n = 6). GAPDH is used as loading control. *Tfrc* expression is depicted as bars representing mean±SEM, n = 6; shown as black dots. p value is determined by the Welch’s one-way ANOVA as differences between untreated group, and Baf- and Sal-treatments. (H) Representative images of FerroOrange staining of cytoplasmic labile iron pools in control, Baf-treated and Baf-treated cells with iron supplementation. Note the reduced staining in Baf-treated cells relative to other conditions. Images are representative of thirty cells per condition from three independent experiments. (I) mRNA levels of *Tfrc* in control, Baf-treated and Baf-treated cells with iron supplementation (left) or in control, Sal-treated and Sal-treated cell with iron supplementation (right) for 24 hours. Bar graphs depict mean±SEM of four independent experimental measures (shown as black dots). p values represent Welch’s one-way ANOVA with Dunnett’s correction for multiple comparisons, estimated as differences between Baf- or Sal-treated cells and other experimental groups. (J) Whole cell immunoblots of HIF-1α and FTH1 in cells treated with Baf or Baf + iron (left) or with Sal and Sal + iron (n = 4). GAPDH is used as loading control. (K) Transcript levels of HIF-1α target genes in cells treated with Baf or with Baf + iron. The mean±SEM of four biological replicates (black dots) is shown. p values are determined by Welch’s one-way ANOVA with Dunnett’s correction for multiple comparisons (all experimental groups compared to Baf-treated cells). (L) Increased *Tfrc* expression (left) in *Gaa*^−/−^ MEFs (n = 6, depicted as black dots). p value is determined by the unpaired two-tailed t-test with Welch’s correction. Whole cell immunoblots of FTL1 and HIF-1α (right) in *Gaa*^−/−^ MEFs (n = 6). GAPDH is used as loading control. (M) Total, ferrous and ferric iron concentrations in MEFs prepared from *Gaa*^−/−^ and their wild type littermate controls. Results are summarized as mean±SEM of experimental measures shown as black dots. Differences between means shown as actual p values are determined by the unpaired two-tailed t-test. See also Figure S1.

The major coordinator of the cellular reponse to hypoxia is the transcription factor hypoxia-inducible factor-1α (Majmundar et al., 2010), which is included in the list of TFs responding to bafilomycin. Furthermore, HIF-2α (EPAS1), whose activation mechanism is similar to HIF-1α, is also part of the TF list. The HIF transcription factors function as heterodimers of a regulated α-subunit and the constitutive β subunit (Majmundar et al., 2010). The α subunits are regulated post-translationally by the prolyl hydroxylases. These enzymes are dioxygenases of the α-ketoglutarate-dependent superfamily, and hydroxylate HIF-1α and HIF-2α in the presence of O_2_, Fe^2+^ and α-ketoglutarate (Majmundar et al., 2010). Hydroxylated HIF-1α and HIF-2α are then recognized by the ubiquitin ligase VHL and targeted to the proteasome for degradation. Notably, the protein levels of HIF-1α were increased in bafilomycin (Baf)-treated fibroblasts (Figure 1D). A similar result was obtained in cells treated with a different inhibitor of the lysosomal v-ATPase, saliphenylhalamide (henceforth, saliphe [or Sal in figures]) (Figure 1D). Accumulation of HIF-1α can be caused by decreased function of the prolyl hydroxylases or decreased levels of O_2_ or VHL (Raimundo et al., 2011). Because the cells are maintained under normoxic conditions, the lack of O_2_ is not a factor. We therefore tested if VHL, α-ketoglutarate or Fe^2+^ could explain the accumulation of HIF-1α and HIF-2α. The protein levels of VHL were not changed in bafilomycin-treated fibroblasts (Figure 1D). To test if the reason why HIF-1α was accumulating upon v-ATPase inhibition was due to decreased levels of α-ketoglutarate, we added a cell-permeant form of α-ketoglutarate to the bafilomycin-treated fibroblasts, and found no decrease in the accumulation of HIF-1α (Supplemental Figure S1A). Then, we tested if there were any perturbations in Fe levels. It is known that iron chelation results in HIF-1α accumulation (Wang and Semenza, 1993). As positive control for the HIF-1α protein accumulation, we treated cells with an iron chelator, deferoxamine (Dfo), which resulted in the expected robust increase in HIF-1α protein levels (Figure 1D). We therefore assessed the total, Fe^2+^ and Fe^3+^ levels in bafilomycin-treated and control fibroblasts, and found a robust decrease in total, Fe^2+^ and Fe^3+^ (Figure 1E). The hydroxylation of HIF-1α, which leads to its ubiquitination and degradation, is Fe^2+^-depedent, and therefore lower Fe^2+^ levels are in agreement with the accumulation of HIF-1α. Altogether, these results suggest that perturbation of Fe homeostasis by v-ATPase inhibition triggers the pseudo-hypoxia HIF-mediated response.

### Lysosomal Fe^2+^ efflux regulates HIF-mediated response

The cytoplasmic concentration of Fe^2+^ is controlled by the iron regulatory protein 1 (IRP1), IRP2 and the ferritinophagy receptor NCOA4 (Mancias et al., 2014; Rouault, 2015). When the levels of cytoplasmic Fe^2+^ are down, IRP activity mediates reduced expression of the iron storage protein ferritin and an increase in transferrin receptor leading to extracellular (transferrin-bound) iron uptake (Rouault, 2015). In parallel, intracellular iron is mobilized by autophagy of ferritin (ferritinophagy) (Mancias et al., 2014) and mitochondria (mitophagy) (Allen et al., 2013; Schiavi et al., 2015), which deliver iron directly to the lysosomes (via autophagosomes), while transferrin receptor uptake first releases iron in the endosomes, which is then delivered by kiss-and-run to mitochondria (Hamdi et al., 2016) or released to the cytoplasm. In the endosomes/lysosomes, Fe^3+^ is reduced to Fe^2+^ by the enzyme STEAP3, and Fe^2+^ (but not Fe^3+^) is released to the cytoplasm by DMT1 (SLC11A2) or MCOLN1 (Figure 1F) (Dong et al., 2008; Touret et al., 2003).

We first tested if the lower cellular Fe^2+^ levels when the v-ATPase is inhibited can affect iron homeostasis. We monitored cellular iron homeostasis by measuring the protein levels of ferritin light (FTL1) and heavy chain (FTH1) subunits and the transcript levels of transferrin receptor (TFRC). The lysosomal chelator Dfo, was used as positive control given that it retains iron in the lysosome and thus impedes its release to the cytoplasm, thus triggering functional iron deficiency (Doulias et al., 2003; Kurz et al., 2006). As TFRC mRNA has a 3’ iron responsive element, its transcript levels are expected to increase upon iron deficiency. We observed in cells treated with the v-ATPase inhibitors, bafilomycin and saliphe, increased *Tfrc* transcript levels and decreased protein amounts of FTL1 and FTH1 (Figure 1G). As expected, Dfo treatment resulted in increased *Tfrc* transcript (Supplementary Figure S1B) and decreased protein levels of the ferritin chains FTL1 and FTH1 (Figure 1G). Importantly, the treatment with Dfo does not impact lysosomal function, as assessed by the protein levels of autophagy markers SQSTM1 and LC3B-II (Supplementary Figure S1C). Furthermore, we stained cells with the probe FerroOrange, which reacts specifically with Fe^2+^ but not Fe^3+^, and found that the signal intensity of FerroOrange was sharply decreased in bafilomycin- or saliphe-treated cells (Figure 1H).

The iron-deficiency response caused by v-ATPase inhibition is likely due to impairment of the iron homeostasis in the endolysosomal system. To test this premise, we supplemented the growth medium with Fe-citrate, which allows iron to be imported through transporters in the plasma membrane (Ofer et al., 1981), thus bypassing the endolysosomal pathway. When cells are treated simultaneously with v-ATPase inhibitors and Fe-citrate, the iron-deficiency response is deactivated, as assessed by the transcript levels of *Tfrc* (Figure 1I). To ensure that his effect was specifically due to Fe, Na-citrate was used as control. This result underscores that the iron homeostatic response is regulated by the cytoplasmic iron levels (Rouault, 2015), and that retention of iron in the lysosome triggers cytoplasmic iron deficiency, which can be resolved by iron import independently of the endolysosomal pathway.

To directly test if the accumulation of HIF-1α upon v-ATPase inhibition was due to the perturbation in Fe homeostasis, we supplemented the growth medium of the bafilomycin- or saliphe-treated fibroblasts with Fe-citrate, which restored HIF-1α to the barely detectable amounts observed in control cells (Figure 1J). The striking normalization of ferritin levels upon Fe-citrate supplementation of bafilomycin-treated cells further underscores that Fe-citrate is being taken up by the cells and resolves the iron deficiency (Figure 1J).

To further probe the impact of HIF-1α accumulation upon v-ATPase inhibition, we tested if the transcriptional targets of HIF-1α were affected. We observed that the transcript levels of HIF-1α targets are robustly induced in bafilomycin- and saliphe-treated cells, and are restored to basal levels when the cells are co-treated with v-ATPase inhibitors and Fe-citrate (Figure 1K and Supplementary Figure S1D). We then sought to test whether genetic defects that impair lysosomal acidification also impact HIF accumulation. Therefore, we used fibroblasts derived from mice lacking acid α-glucosidase (*Gaa*, mice referred henceforth as *Gaa*-KO), a lysosomal enzyme mutated in Pompe’s disease, in which a large portion of lysosomes is not able to acidify (Fukuda et al., 2006). We observed that the *Gaa*-KO fibroblasts present iron deficiency as assessed by increased *Tfrc* transcript level, and decreased FTL1 levels (Figure 1L). Notably, these fibroblasts also present HIF-1α accumulation (Figure 1L). Furthermore, the *Gaa*-KO fibroblasts show a decrease in Fe^2+^, in agreement with the iron deficiency response and the accumulation of HIF-1α (Figure 1M). The increase in Fe^3+^ suggests a defect in the endolysosomal reduction of Fe^3+^ to Fe^2+^.

Altogether, these results show that v-ATPase inhibition results in decreased cytoplasmic Fe^2+^, which impairs the hydroxylation-mediated degradation of HIF-1α and culminates in the accumulation and activation of this transcription factor.

### Mitochondrial biogenesis and function require iron

Having shown that endolysosomal acidification is required for Fe^2+^ release from endo/lysosomes to the cytoplasm, we sought to test if mitochondrial iron content was also affected. We loaded the cells with the probe Mito-FerroGreen, which localizes to mitochondria and reacts specifically with Fe^2+^. The cells treated with bafilomycin showed a robust decrease in Mito-FerroGreen signal (Figure 2A). This effect was ablated in the presence of Fe-citrate (Figure 2A). This result shows that supply of iron to mitochondria is also impaired when the v-ATPase is inhibited, and that mitochondria seem to be able to uptake iron from cytoplasm when the endolysosomal pathway is ineffective.

**Figure 2.**
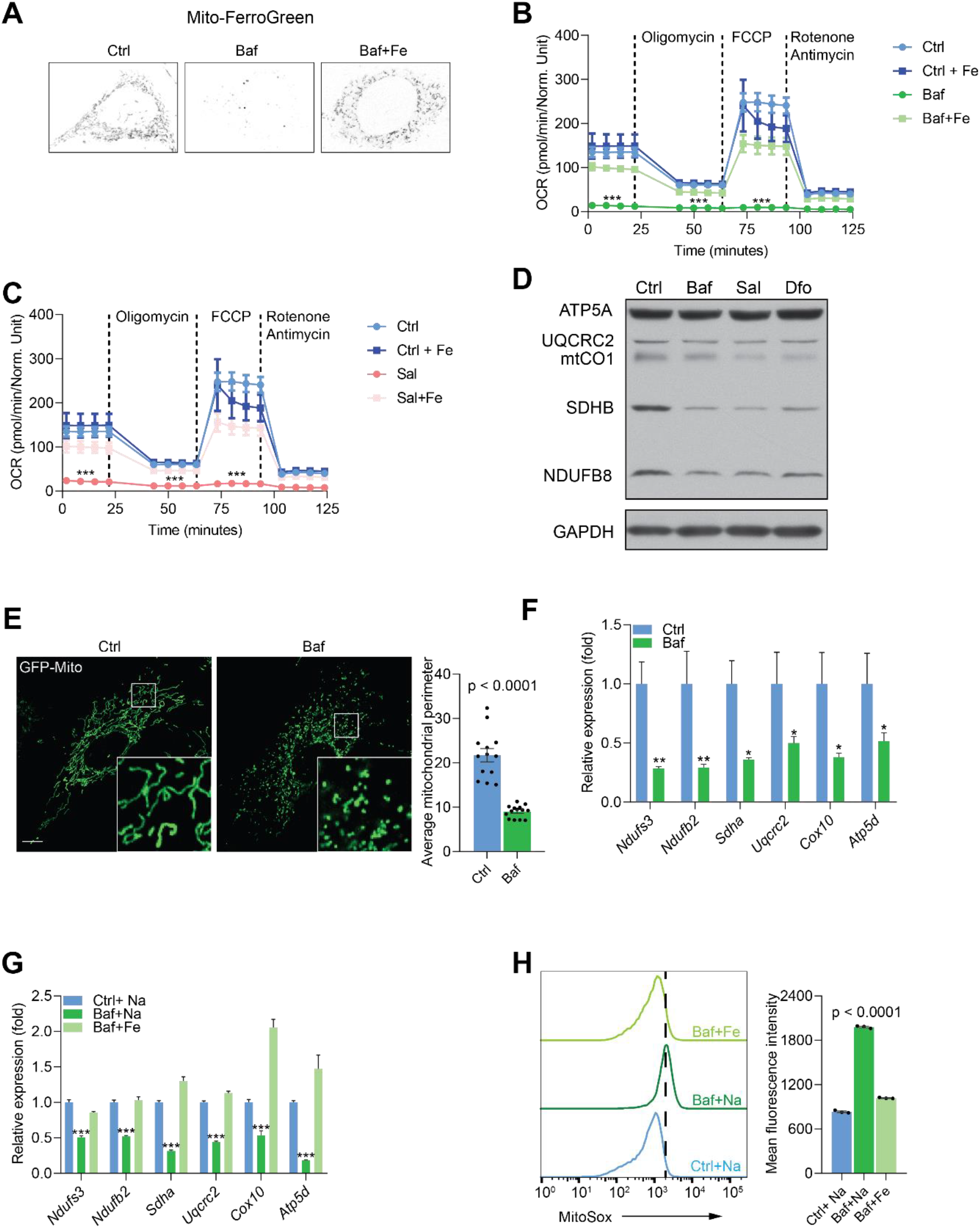
Mitochondrial biogenesis and function are dependent on endolysosomal iron supply. (A) Representative image Mito-FerroGreen staining of mitochondrial labile iron pools in control, Baf- and Baf + Fe-treated cells. Note the barely detectable labile iron levels in mitochondria. Images represent results from thirty cells each per experimental condition from triplicate experiments. (B-C) Mitochondrial oxygen consumption rates (OCR) in Baf- and Baf-treated cells with iron supplementation (B) or in Sal- and Sal-treated cells with iron supplementation (C). Results represent mean±SEM, three independent experimental replicates. Each experimental replicate is calculated from the average of 8 technical replicates. ***p < 0.001, Welch’s one-way ANOVA with Dunnett’s correction for multiple comparison (all experimental groups compared to Baf- or Sal-treated cells). (D) Whole cell Western blot of ATP5A, UQCRC2, mtCO1, SDHB and NDUFSB in control, Baf- and Sal-treated cells (n = 6). Dfo is used as positive control for iron deficiency. Note the reduction of mitochondrial proteins in treated cells relative to controls. GAPDH is used as loading control. (E) Representative images of cells transfected with GFP-Mito and treated with Baf for 24 hours to show the mitochondrial network. Note the prevalent mitochondrial fragmentation in Baf-treated cells. Number of cells is shown as black dots per condition in bars representing mean±SEM of mitochondrial perimeter from three independent experiments. p value is estimated by the Mann-Whitney U-test. Scale bar, 10µM. (F) Transcript levels of nuclear-encoded mitochondrial genes in Baf-treated cells. Bars depict mean±SEM of six independent experimental measures. *p < 0.05; **p <0.01; ***p < 0.001, unpaired two-tailed t-test with Welch’s correction. (G) Trancript levels of nuclear-encoded mitochondrial genes in Baf-treated cells and Baf-treated cells with iron supplementation. Results are shown as mean±SEM, n = 8. *p < 0.05; **p <0.01; ***p < 0.001, Welch’s one-way ANOVA with multiple test correction made by the Dunnett’s method. All comparisons were made between Baf-treated cell and other experimental conditions. (H) Mitochondrial superoxide levels in Baf- and Baf-treated cell with iron supplementation. Differences are depicted as mean fluorescent intensities of the superoxide-sensitive and mitochondrial-targeted dye Mitosox. Error bars represent SEM of three independent experimental measures (black dots). p value is Welch’s one-way ANOVA with Dunnett’s correction for multiple corrections (all conditions compared to Baf-treated cells). See also Figure S2.

Mitochondria are a key component of Fe-S cluster synthesis, which is essential for the proper function of the respiratory chain. To assess if mitochondrial function was impacted by the decrease in Fe^2+^, we performed real-time respirometry, which monitors O_2_ consumption in real-time as a proxy for mitochondrial respiratory chain activity. We observed that cells treated with either v-ATPase inhibitor bafilomycin or saliphe showed a robust decrease in mitochondrial O_2_ consumption (Supplementary Figure S2A). Interestingly, treatment of cells with the lysosomal iron chelator Dfo resulted in virtual absence of mitochondrial respiratory chain activity (Supplementary Figure S2B). Notably, supplementation of the medium with Fe-citrate was sufficient to return mitochondrial respiratory chain activity to normal levels in cells treated with bafilomycin (Figure 2B), saliphe (Figure 2C) or with Dfo (Supplementary Figure S2C).

We then explored what may be driving the decrease in mitochondrial respiration. Multiple causes are possible, including a decrease in the mass or the quality of the mitochondrial network, or a metabolic shift that shunts pyruvate away from aerobic metabolism. First, we assessed the protein levels of respiratory chain subunits in whole cell extracts by Western blot, and observed that they are decreased when v-ATPase is inhibited (Figure 2D). A similar result is observed in cells treated with the iron chelator Dfo (Figure 2D). To assess mitochondrial mass, we measured the protein levels of the proteins VDAC1 and Tom20, which are highly abundant in the outer mitochondrial membrane. We found a decrease in VDAC1 and Tom20 levels in v-ATPase-inhibited cells (Supplementary Figure S2D). Next, we tested if the mitochondrial network was affected by v-ATPase inhibition, and observed a robust phenotype of mitochondrial fragmentation and swelling (Figure 2E).

Because autophagy is impaired in cells treated with v-ATPase inhibitors (Supplementary Figure S1D), increased mitophagy cannot explain lower mitochondrial mass. Therefore, we tested if the transcriptional program of mitochondrial biogenesis was affected. We measured the transcript levels of several mitochondrial respiratory chain subunits (all encoded in the nuclear DNA), and found that they all presented a robust down-regulation when treated with bafilomycin (Figure 2F) or with saliphe (Supplementary Figure S2E). A similar effect was observed when the cells were treated with Dfo, as previously shown by Pagliarini and colleagues (Rensvold et al., 2013). Notably, supplementation of the cell medium with Fe-citrate returned the transcript levels of the mitochondrial respiratory chain subunits to control levels in bafilomycin (Figure 2G) or saliphe treatment (Supplementary Figure S2F). The protein levels of several mitochondrial subunits also show a recovery to control levels when Fe-citrate is given together with bafilomycin (Supplementary Figure S2G).

Finally, we also assessed the efficiency of electron transfer along the mitochondrial respiratory chain by estimating the levels of superoxide (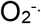, the product of a single-electron reduction of O_2_), which is a byproduct of inefficient electron transfer along the chain. Using the mitochondria-targeted superoxide-sensitive dye MitoSox, we observed a sharp increase in MitoSox levels in bafilomycin-treated cells, almost completely recovered in the cells that also received Fe-citrate (Figure 2H). We probed the levels of pro-oxidant molecules (superoxide, hydroxyl radical, among others) using the probe dichlorofluoresceine (DCF), which reacts with many reactive oxygen species (ROS). We observed a sharp increase in DCF intensity in bafilomycin-treated cells, which was normalized by co-treatment with Fe-citrate (Supplementary Figure S2H). A similar pattern is observed with the dye BODIPY C11, which reports the levels of oxidized membrane phospholipids (Supplementary Figure S2I). The increased levels of mitochondrial superoxide, ROS and lipid peroxidation were recapitulated in saliphe-treated cells and rescued following co-treatment with Fe-citrate (data not shown). Overall, these results show that mitochondrial biogenesis and function require iron, made available to the cytoplasm by the endolysosomal system or via direct transport through the plasma membrane (Fe-citrate supplementation). In the absence of iron, mitochondrial function is impaired and generates more superoxide, promoting an oxidative shift in the cellular redox environment.

### Lysosomal Fe^2+^ efflux is required for cell proliferation in a mitochondrial respiratory chain-dependent manner

Given the importance of iron to many cellular processes, we tested whether iron-deficiency induced by v-ATPase inhibition could affect cell proliferation. We observed that cells treated with vehicle control proliferate, while both bafilomycin (Figure 3A) and saliphe treatments (Figure 3B) result not only in halting of cell proliferation but also in cell death (the number of cells decreases during the treatment). The treatment with saliphe has slightly lower impact on cell death than with bafilomycin (Figure 3B). Remarkably, the supplementation of the medium with Fe-citrate during v-ATPase inhibition restores cellular proliferation (Figure 3C). Similar results were observed in cells treated with the iron chelator Dfo (Supplementary Figure S3A-B).

**Figure 3.**
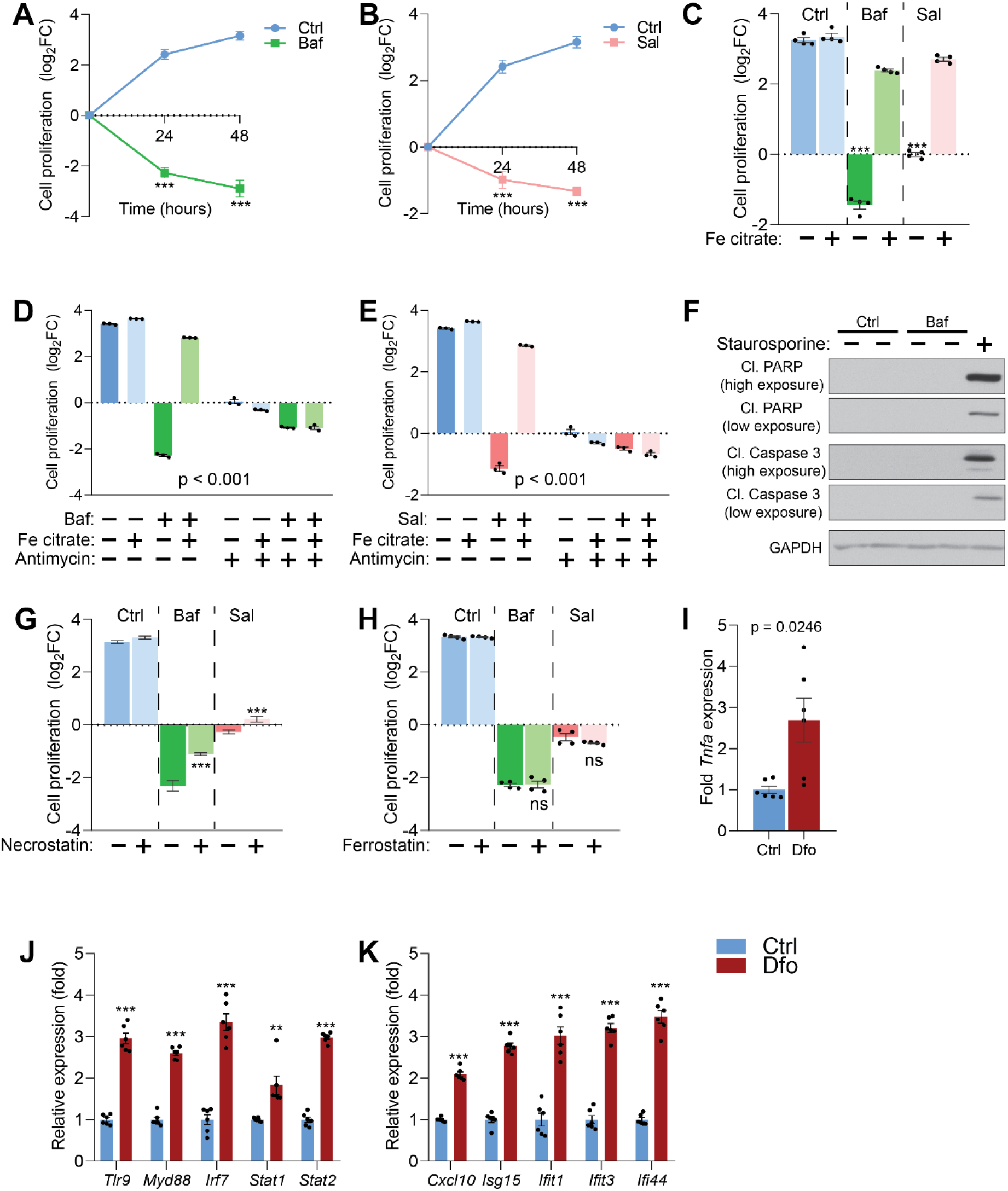
Lysosomal iron efflux abrogates necrotic cell death. (A-B) Baf-treated (A) and Sal-treated (B) cells display about 4-to 8-fold increase in cell death with time of treatment, relative to untreated cells, which show progressive cell proliferation. Results are presented as mean±SEM, n = 4 experimental replicates with each experimental replicate being the average of technical triplicates. ***p < 0.001, unpaired two-tailed t-test with Welch’s correction (C) Iron supplementation rescues cell death upon treatment with Baf or Sal. Bars and error bars represent mean±SEM of log2 fold change in cell number of experimental conditions. Black dots indicate individual experimental measures. *** p < 0.001, Welch’s one-way ANOVA with Dunnett’s correction for multiple comparisons. All conditions compared to cells treated with either Baf or Sal alone. (D-E) 20µM Antimycin treatment for 48 hours abolishes the rescue of cell death following iron supplementation in Baf-treated (C) or Sal-treated (D) cells. Bar graphs represent mean±SEM of three independent experimental measures (black dots). *** p < 0.001, Two-way ANOVA with Sidak correction for multiple comparisons. (F) Whole cell immunoblot of cleaved PARP and cleaved caspase-3 levels shows that Baf-induced cell death is non-apoptotic (n = 3). Staurosporine (1µM treatment for 4 hours) is used as positive control for caspase-3 dependent apoptotic cell death. GAPDH is used as loading control. (G-H) Necrostatin treatment (40µM) for 48 hours partially rescues cell death induced by Baf- or Sal-treatment in cells (G). 5µM Ferrostatin treatment for 48 hours has no observed effect on Baf or Sal-induced cell death in cells. Results show mean±SEM, n = 6 (H) and n = 4 (black dots in G). *** p < 0.001; ns p >0.05, Welch’s one-way ANOVA with Dunnett’s correction for multiple comparisons to treatment with Baf or Sal alone. (I-K) Deferoxamine treatment in neurons results in more than 2-fold increase in the expression of *Tnfa* (I), increased expression of regulators of interferon gene expression (J) and increased transcript levels of interferon-stimulated genes (K). Results show mean±SEM of six independent experiments. **p<0.01; ***p <0.001, unpaired two-tailed t-test with Welch’s correction. See also Figure S3 and Tables S1, S2 and S3.

Because iron supplementation in the presence of v-ATPase inhibitors also improved mitochondrial function, we tested if the mitochondrial respiratory chain activity was necessary for the recovery in cell viability and proliferation. The addition of the mitochondrial respiratory chain complex III inhibitor antimycin A to bafilomycin-treated iron-supplemented cells ablates the viability recovery conferred by the iron supplementation (Figure 3D). A similar result is observed in saliphe-treated (Figure 3E) and in Dfo-treated cells (Supplementary Figure S3C).

Next, we sought to characterize the cell death phenotype in response to v-ATPase inhibition. We first looked for apoptotic markers such as cleaved (active) caspase 3 and cleaved PARP (caspase 3 substrate). We could not detect either of them in bafilomycin-(Figure 4F) or saliphe-treated cells (Supplementary Figure S3D), despite they were readily detectable in cells treated with the apoptosis inducer staurosporine. Accordingly, pan-caspase inhibitor ZVAD had no attenuating effect on cell death in bafilomycin- or saliphe-treated cells (Supplementary Figure S3E), further supporting that the cell death in these conditions is not apoptotic. We then tested if the cell death induced by v-ATPase inhibition was sensitive to necrostatin (inhibitor of necrosis) or to ferrostatin (inhibitor of ferroptosis). Necrostatin robustly reduced the cell death caused by v-ATPase inhibition (Figure 3G). Ferrostatin had no effect on v-ATPase inhibition-induced cell death (Figure 3H), despite it effectively inhibited cell death triggered by the ferroptosis inducer erastin (Supplementary Figure S3F). Therefore, necrosis seems to be the main mechanism of cell death in cells subjected to v-ATPase inhibition. In agreement with this finding, the analysis of the three transcriptome datasets of bafilomycin-treated cells that were used in Figure 1 ranks necrosis as the top cell death mechanism (Supplementary tables S1-S3).

### Iron deficiency triggers cell-autonomous inflammatory gene expression

The occurrence of necrosis is often associated with triggering of sterile inflammatory responses (Rock and Kono, 2008). To test if iron deficiency is sufficient to trigger immune responses, we treated mouse primary cortical neurons with Dfo, and monitored the expression of pro-inflammatory cytokines and interferon-stimulated genes, which are typically increased during inflammatory responses. We first observed the expression of tumor necrosis α (*Tnfa*) and found a robust increase in its transcript levels in Dfo-treated primary neurons (Figure 3I). We then measured the transcript levels of innate immune regulators such as *Tlr9*, *Myd88*, *Irf7*, *Stat1* and *Stat2*, which were all robustly up-regulated in Dfo-treated cortical neurons (Figure 3J). Accordingly, the expression of other downstream interferon-stimulated genes such as *Cxcl10*, *Isg15*, *Ifit1*, *Ifit3* and *Ifi44* was also strongly induced in Dfo-treated neurons (Figure 3K). We have not tested the effect of bafilomycin or saliphe treatment in the primary neurons because these v-ATPase inhibitors would not just inhibit lysosomal function but also secretory vesicle (Farsi et al., 2018), and therefore trigger a number of events that are unrelated to iron homeostasis. These results underscore that unavailability of cytoplasmic iron is sufficient to trigger inflammatory signaling and necrotic cell death.

**Table 1.** Relaxation times of water protons and Magnetization Transfer ratios determined by MRI. (related to Figure 4).

|  |  | **6 months** |  |  | **12 months** |  |
| --- | --- | --- | --- | --- | --- | --- |
| Background |  | Wild type | GAA(-/-) |  | Wild type | GAA(-/-) |
| *n* |  | *n* = 8 | *n* = 8 |  | *n* = 5 | *n* = 5 |
| ***T<sub>1</sub>*(s)** |  |  |  |  |  |  |
| **Cerebral Cortex** |  | 1.67 ± 0.07 | 1.67 ± 0.07 |  | 1.65 ± 0.05 | 1.62 ± 0.09 |
| **Striatum** |  | 1.62 ± 0.05 | 1.64 ± 0.07 |  | 1.62 ± 0.03 | 1.58 ± 0.06 |
| **Thalamus** |  | 1.49 ± 0.03 | 1.52 ± 0.09 |  | 1.50 ± 0.04 | 1.46 ± 0.03 |
| ***T<sub>2</sub>*(ms)** |  |  |  |  |  |  |
| **Cerebral Cortex** |  | 40.3 ± 0.5 | 38.3 ± 0.8*** |  | 39.8 ± 0.7 | 37.9 ± 0.5* |
| **Striatum** |  | 39.7 ± 1.0 | 37.3 ± 0.8** |  | 38.5 ± 0.6 | 36.7 ± 0.7* |
| **Thalamus** |  | 38.4 ± 1.0 | 35.9 ± 1.1** |  | 37.2 ± 0.3 | 36.2 ± 0.8 |
| ***Magnetization Transfer Ratio (%)*** |  |  |  |  |  |  |
| **Cerebral Cortex** |  | 7.5 ± 1.7 | 7.0 ± 1.7 |  | 7.8 ± 0.9 | 8.4 ± 0.6 |
| **Striatum** |  | 9.9 ± 0.7 | 9.0 ± 0.8* |  | 9.8 ± 0.6 | 10.0 ± 0.9 |
| **Thalamus** |  | 12.5 ± 1.6 | 11.9 ± 1.9 |  | 12.9 ± 2.0 | 13.0 ± 0.9 |

### Iron deficiency is prevalent in the brain of an *in vivo* model of impaired lysosomal acidification

Given the pro-inflammatory nature of Fe-deficiency caused by lysosomal v-ATPase inhibition, we sought to test the system *in vivo*, using a mouse model lacking *Gaa* (*Gaa*-KO), as described above. Absence of *Gaa* impairs the acidification of a large proportion of lysosomes, and fibroblasts obtained from these mice show iron deficiency and HIF-1α accumulation (see Figure 1L). The *Gaa*-KO mice show a predominantly muscular phenotype, albeit with a later disease onset when compared to human patients with *Gaa* loss-of-function mutations (Raben et al., 1998). Nevertheless, the mice present severe motor symptoms with onset around 14 months, and so we assessed them at earlier time points (6 and 12 months) to avoid this confounding factor. We focused on the brain of the *Gaa*-KO mice, as loss of lysosomal acidification is particularly common in neurodegenerative diseases (Lie and Nixon, 2019). Furthermore, it has been shown that the muscular phenotype of the *Gaa*-KO mice can be rescued by re-expression of *Gaa* in the motor neurons (Lee et al., 2018; Todd et al., 2015; Turner et al., 2016), illustrating that lysosomal function is relevant in neuronal cell populations. First, we tested if the cortex of the *Gaa*-KO mice showed signs of iron deficiency. We observed a robust increase in the transcript levels of *Tfrc* in the cortex of 6- and 12-month-old mice (Figure 4A). Ferritin light chain protein levels were down-regulated both at 6- and 12-months (Figure 4B). Together, these results document a robust iron deficiency response in *Gaa*-KO brain. To further probe the functional consequences of iron-deficiency in the *Gaa*-KO brain, we also tested if HIF-1α was accumulating, by western blot, and found an increase in its levels in *Gaa*-KO cortex both at 6- and 12-months (Figure 4C). Furthermore, mitochondrial iron availability was also affected, as the activity of complex IV, which is dependent on Fe, was robustly decreased in *Gaa*-KO cortex (Figure 4D), while the activity of complex V (loading control) was similar between WT and KO (Figure 4D). The decrease in complex IV activity (cytochrome c oxidase, COX) occurs despite the protein levels of the subunit COX1 were similar between WT and KO both in whole cell extracts (Supplementary Figure S4A) and mitochondrial extracts (Supplementary Figure S4B) from mouse cortices..

We then performed *in vivo* magnetic resonance spectroscopy (MRS) and imaging in the mice at 6 and 12 months of age (Figure 4E), to assess if the perturbations in iron homeostasis were widespread or in discrete brain regions. For this, we analyzed the thalamus, striatum and cerebral cortex, and in all three regions we found a decrease in the T_2_ relaxation time of water protons (Table 1), which is highly correlated with reduced labile iron pool levels (Vymazal et al., 1993). Interestingly, we also observed by MRS, reduced levels of choline-containing compounds, which may be due slow turnover of phosphatidylcholine, further suggesting impaired myelination (Figure 4E and Supplementary table S4). We therefore tested whether myelination (which is also a Fe-dependent process) was affected in the *Gaa*-KO cortex. We observed that the abundance of myelination-related proteins such as proteolipid protein (PLP) and myelin basic protein (MBP) was decreased in *Gaa*-KO cortical homogenates (Supplementary Figure S4C). Altogether, these results show that the *Gaa*-KO brain is iron-deficient, with consequences for HIF signaling, mitochondrial function and myelination.

**Figure 4.**
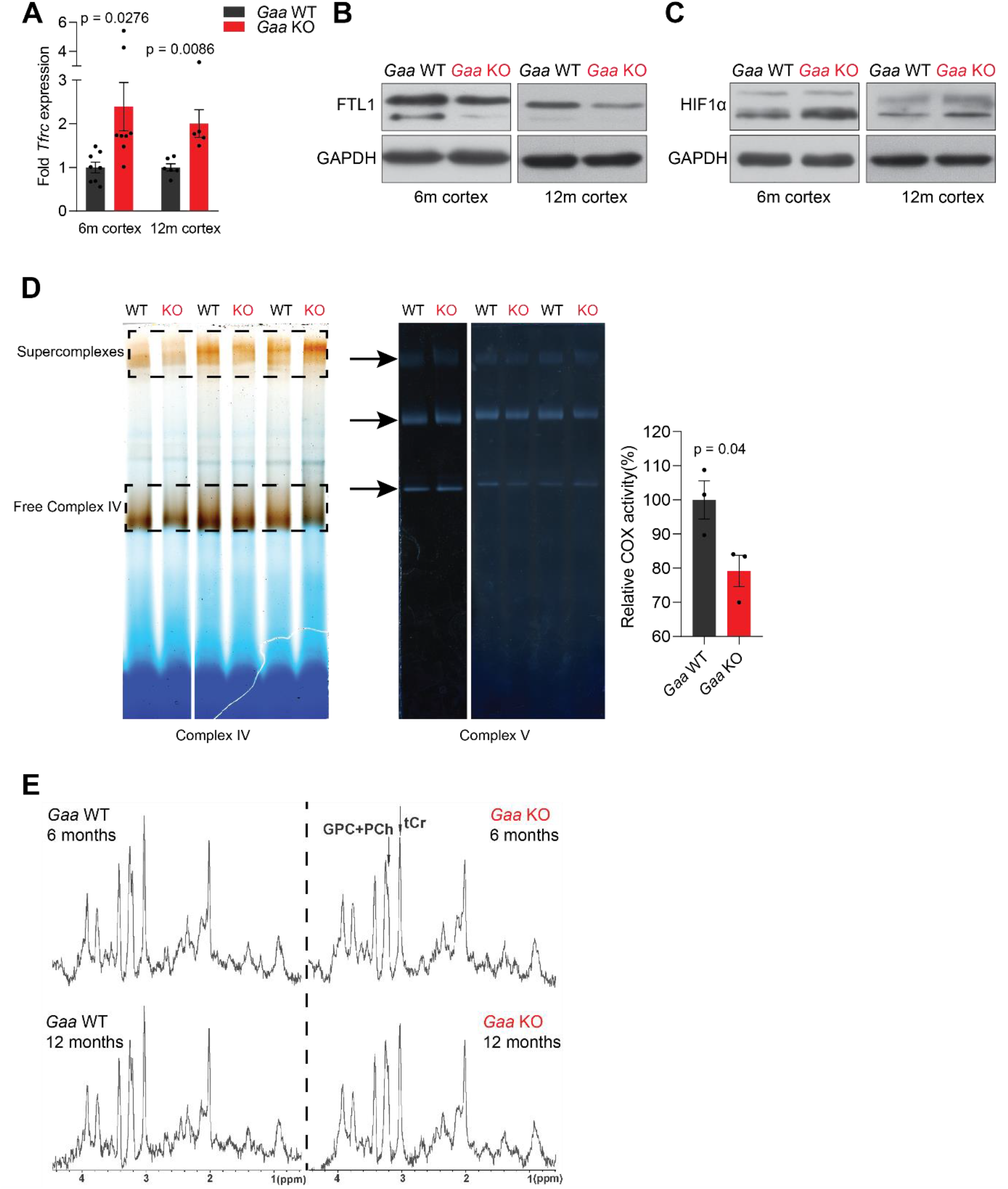
Functional iron deficiency in a mouse model of impaired lysosomal acidification (*Gaa*^−/−^ mouse) (A) *Tfrc* mRNA levels in the cortex of 6 and 12 month old *Gaa*^−/−^ mice and their wild type littermate controls. Black dots represent sample size (n) in each group. p values are unpaired two-tailed t-tests with Welch’s correction. Bars are depicted as mean±SEM. (B) Western blot of FTL1 and GAPDH as loading control from cortical tissue lysates of 6 and 12 month old *Gaa*^+/+^ and *Gaa*^−/−^ mice (n = 4 mice per group). (C) Whole tissue immunoblots showing increased HIF-1α levels with GAPDH as loading control in wild type and *Gaa*^−/−^ mice at 6 and 12 months of age (n = 4 animals per group). (D) Activity staining of native respiratory chain complex IV of mitochondria purified from cortices of 6-month old mice (n = 3; depicted as black dots). Complex V activity staining is used as loading control. Arrows mark complex V activity staining. Top arrow, dimers; middle arrow, monomers; bottom arrow, unassembled F1. The difference between complex IV activities (mean±SEM) of groups is determined by the unpaired two-tailed t-test with Welch’s correction. (E) MR spectra reveal significant changes in metabolite concentrations in *Gaa*^−/−^ mice *in vivo*. MRS (STEAM, TR/TE/TM = 6000/10/10 ms, 256 averages, (2.0 mm)^3^ volume-of-interest centered on the striatum) of 6-month old (upper row) and 12-month old (lower row) wild type (left column) and *Gaa*^−/−^ mice (right column) *in vivo*, as summarized in Table S4, processed with a 1-Hz line broadening. GPC+PCh = Choline-containing compounds, tCr = total creatine, ↑ = significant signal intensity increases, ↓ = significant signal intensity decreases. *p < 0.05; **p < 0.01, Mann-Whitney’s U-test. See also Figure S4, Table 1 and Table S4.

### *Gaa*-KO brain shows prevalent inflammation at very early presymptomatic stages

Having shown that the *Gaa*-KO brain presents functional iron deficiency akin to what was observed in cultured *Gaa*-KO cells, as well as in cells treated with v-ATPase inhibitors, we sought to determine if we could detect inflammatory signatures in the *Gaa*-KO mouse cortex. In order to have an unbiased approach, we performed RNA sequencing of WT (n = 5) and *Gaa*-KO (n = 5) 12 month-old cortices. We identified 1779 differentially expressed genes (adjusted p value < 0.05; fold change > 2.0), of which 996 were up-regulated and 783 down-regulated in *Gaa*-KO cortices.The differential gene list fully segregates WT and KO samples in hierarchical clustering (Figure 5A). The most significantly changed transcripts are *Gaa* (as expected) and several proteins related to immune responses, particularly complement activation, macrophage infiltration and genes induced by interferon signaling (Figure 5B). Accordingly, several pathways related to inflammation were found enriched in the *Gaa*-KO dataset (Figure 5C). We then determined which upstream regulators (specifically, transcription factors) were affected in the *Gaa*-KO cortex, and found that the interferon regulatory factors *Irf7* and *Irf3* were predicted as the two most active transcription factors (Figure 5D). Thus, we focused on the target genes of *Irf7*, and using the *Gaa*-KO cortex transcriptome data we observed that 17 out of 20 *Irf7* targets in the *Gaa*-KO brain are robustly up-regulated (Figure 5E). Altogether, these results show in an unbiased manner that the 12-month old cortex of the *Gaa*-KO mouse shows a robust inflammatory signature involving *Irf7* signaling.

**Figure 5.**
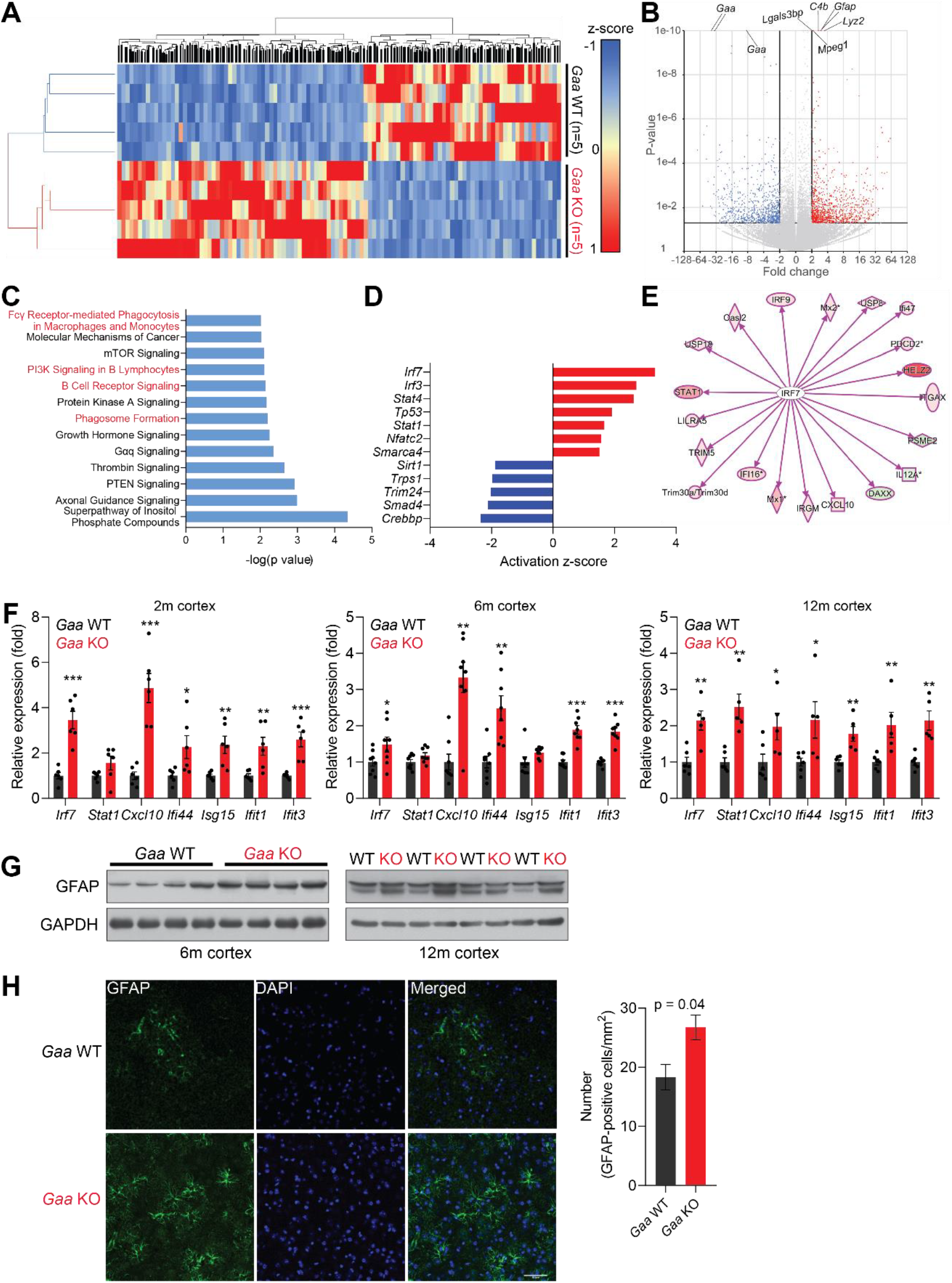
Iron deficiency is linked to induction of innate immunity in the CNS of *Gaa*^−/−^ mice. (A) Hierarchical clustering of the brain samples of *Gaa*^−/−^- and WT mice at 12 months of age. (B) Volcano plot of the transcripts detected in the RNAseq analysis, highlighting those that were considered as differentially expressed genes (adjusted p value < 0.05, and fold-change > 2). Some of the transcripts with the lowest p value between WT and KO are highlighted in the plot. (C) Results of the pathway analysis showing the pathways most enriched in the differentially expressed gene list of the *Gaa*-KO/WT cortex dataset. The pathways related to inflammation are highlighted in red. (D) Transcription factors predicted to be significantly active or repressed in the cortex of 12-month old *Gaa*^−/−^ mice compared to WT. The z-score is represented (positive z-score, predicted activation; negative z-score, predicted repression). (E) Transcript levels of I*rf7* targets recognized in the *Gaa*^−/−^ dataset by Ingenuity Pathway Analysis. The targets are color-coded (red indicates higher expression in *Gaa*^−/−^ mice, green represents lower expression), showing that the vast majority is up-regulated, in agreement with the predicted activation of *Irf7*. (F) Transcript levels of interferon-stimulated genes and their regulators in the cortices of 2-month, 6-month and 12-month old *Gaa*^−/−^ mice and their wild type littermate controls. Number of mice per group is displayed as black dots in bars, which represent mean±SEM. *p <0.05; **p<0.01; ***p <0.001, unpaired two-tailed t-test with Welch’s correction. (G) Whole cortical tissue immunoblots of increased GFAP levels and GAPDH as loading control in 6- and 12-month old wild type and *Gaa*^−/−^ mice (n = 4 mice per group). (H) Representative images of GFAP (green) in the cortex of 6-month old *Gaa*^−/−^ mice and their wild type littermate controls. Scale bars, 20µM. Quantifications on the right show the increased number of GFAP-positive cells in the cortices of *Gaa*^−/−^ mice (n = 3 mice per group) depicted as bars denoting mean±SEM. p value is estimated from an unpaired two- tailed t-test with Welch’s correction.

Next, we tested if the inflammatory signature in the *Gaa*-KO cortex can be detected at earlier ages. We measured by qPCR the transcript levels of several transcripts regulated by *Irf7*, which were up-regulated in the *Gaa*-KO brain at 2, 6 and 12 months of age (Figure 5E). The exception was *Sta1*, which was up-regulated only at 12-months. Glial activation is a typical marker of brain inflammation, and can be assessed by the levels of the microglia- and astrocyte-enriched protein GFAP (which is highly up-regulated at transcript level in the RNA sequencing dataset, see Figure 5B). Therefore, we searched for markers of glial activation in the *Gaa*-KO cortex. We observed an increase in GFAP protein levels in GAA-KO cortex at 6 and 12 months (Figure 5G), by western blot. Staining of tissue sections against GFAP protein in the 6-month GAA-KO cortex yielded similar results (Figure 5G).

Altogether, these results show that the brain of GAA-KO exhibits robust inflammatory and interferon signatures and enrichment of innate immune cells detectable already at 2 months of age.

### Iron deficiency perturbs mtDNA homeostasis *in vivo*

Mitochondrial malfunction has been shown to be a contributor to inflammatory reactions (West and Shadel, 2017). In particular, imbalance in mtDNA homeostasis is a robust trigger of type I interferon responses (West et al., 2015). Therefore, we sought to test if iron deficiency, which we showed to be sufficient to trigger inflammation in cultured neurons (see Figures 3I-K), was associated with mtDNA perturbation *in vivo*. First, we measured mtDNA copy number in the *Gaa*-KO cortex. We observed a reduction in mtDNA of about 20% in newborn mice, 25% at 2 months, and a robust decrease of ~50-60% in 6- and 12-month old *Gaa*-KO cortices (Figure 6A). Notably, the protein levels of TFAM, a key protein for mtDNA replication, transcription and maintenance, were reduced by ~50% in the *Gaa*-KO cortex (Figure 6B). This decrease is akin to the loss of TFAM and mtDNA observed in *Tfam* heterozygous knockout mice, in which mtDNA instability was shown to trigger interferon signaling (West et al., 2015). The decrease in TFAM protein levels cannot be explained only by decreased transcription, since *Tfam* transcript levels are only 20% down in the *Gaa*-KO brain (Supplementary Figure S5A).

**Figure 6.**
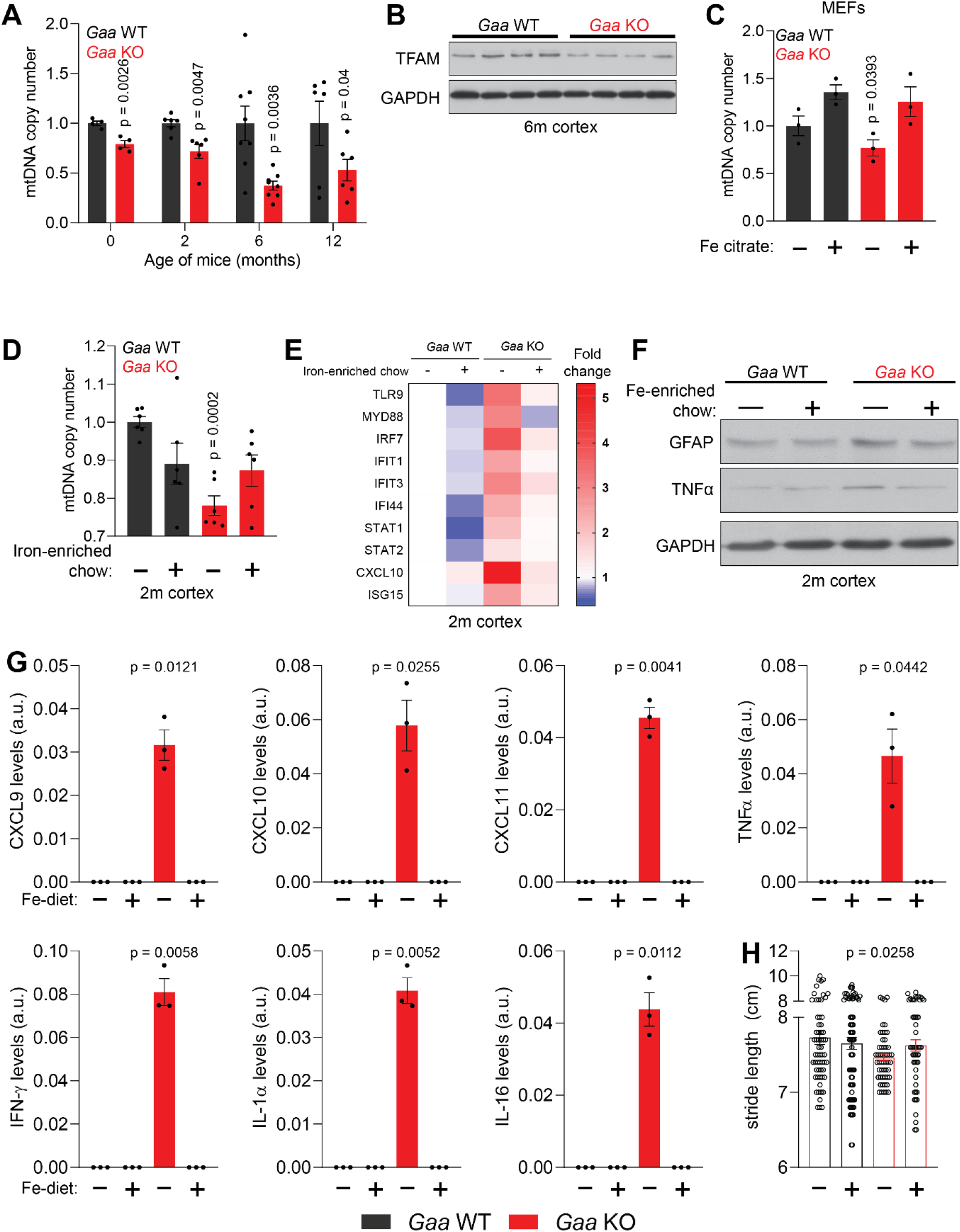
Iron deficiency impairs mtDNA homeostasis in *Gaa*^−/−^ mice. (A) Age-dependent decline in relative mtDNA copy number levels in the cortex of *Gaa*^−/−^ mice from 0 to 12 months of age. Number of mice per group is depicted as dots in bars which denote mean±SEM. *p < 0.05; **p < 0.01; ***p < 0.001, unpaired two-tailed t-test with Welch’s correction. (B) Western blot showing reduced TFAM and GAPDH as loading control in the cortex of *Gaa*^−/−^ mice and their wild type littermate controls (n = 8 mice per group). (C-D) Iron supplementation rescues mtDNA copy number defects in *Gaa*^−/−^ MEFs (C) and in the cortex of 2-month old *Gaa*^−/−^ mice (D). Results are summarized as mean±SEM of n = 3 independent measures in *Gaa*^−/−^ MEFs and n = 6 mice per group indicated as black dots for 2-month old *Gaa*^−/−^ mice. *p < 0.05; ***p < 0.001, Welch’s one-way ANOVA test with Dunnett’s correction for multiple comparisons. All conditions compared to the untreated *Gaa*^−/−^ MEFs (C) or *Gaa*^−/−^ mice fed standard chow (D). (E) Iron supplementation as iron-enriched chow dampens the increased expression of innate immune response genes in *Gaa*^−/−^ mice at 2 months of age. This result is depicted as a Heatmap for n = 6 mice per group, in which blue denotes decreased expression of interferon-stimulated genes relative white (no change or baseline) and red denotes increased expression. Note the dampening of the hot red in *Gaa*^−/−^ on standard chow relative the cohort of *Gaa*^−/−^ mice on iron-enriched chow. (F) Immunoblot of GFAP and TNFα in the cortices of *Gaa*^−/−^ mice and their wild type littermate controls fed standard or iron-enriched chow for 2 months (n = 3). Note the reduction in GFAP and TNFα protein amounts following iron supplementation in *Gaa*^−/−^ mice. GAPDH is used as loading control (G) Iron-enriched diet restores increased levels of cytokine and chemokines to control levels in the cortices of 2-month old *Gaa*^−/−^ mice. Cytokines are increased in *Gaa*^−/−^ mice fed standard chow, and are barely detectable in *Gaa*^−/−^ mice fed iron-enriched diet or their wild type littermate controls. Levels of cytokines and chemokines are shown as mean±SEM (n = 3 mice per group, black dots) and p values represent Brown-Forsythe’s one-way ANOVA tests estimated as differences between *Gaa*^−/−^ mice on standard chow and all other experimental groups. (H) Mouse gait analysis shows rescue of stride length following 2 months of iron supplementation in *Gaa*^−/−^ mice. Data is presented as mean±SEM. Each experimental data point (hollow circles) represent stride length measured on all four limbs of each animal. 13-19 animals per group were used. p value is determined by the Welch’s one-way ANOVA followed by Dunnett’s correction for multiple comparisons of all groups to *Gaa*^−/−^ mice fed standard diet. See also Figure S5.

We then turned to clonal cells to test if the loss of mtDNA observed *in vivo* could be triggered in cells by iron deficiency. We first measured mtDNA copy number in WT MEF treated with the lysosomal iron chelator Dfo, and observed a reduction of ~40% in mtDNA (Supplementary Figure S5B). A similar result was obtained by Dfo treatment of primary cortical neurons (Supplementary Figure S5C). To test the effect of iron deficiency on mtDNA in the absence of lysosomal perturbations, we used MEFs obtained from mice lacking the iron regulatory proteins IRP1 and IRP2, in which the expression of heavy and light ferritin chains is robustly up-regulated, and for that reason are functionally iron deficient (all available iron is stored in the ferritin oligomers, and thus biologically inactive). The mtDNA copy number in the *Irp1*^−/−^ and *Irp2*^−/−^ KO MEFs was ~40% reduced compared to WT (Supplementary Figure S5D). We then tested whether impaired lysosomal acidification caused by genetic mutations also resulted in decreased mtDNA copy number. To this end, we used MEFs obtained from *Gaa*-KO and respective WT littermates, and observed a loss of ~40% mtDNA in *Gaa*-KO (Supplementary Figure S5E). Notably, the protein levels of TFAM were also decreased in *Gaa*-KO MEFs (Supplementary Figure S5F).

mtDNA synthesis relies on dNTPs imported from the cytoplasmic dNTP pool (Copeland, 2012), which is high in proliferating cells but decreases sharply in post-mitotic cells and tissues. Ribonucleotide reductase is a key enzyme for the reduction of NTPs to dNTPs, and its active site is dependent on iron (Puig et al., 2017). Thus, the iron deficiency observed in *Gaa*-KO cells or Dfo treated cells, could result in a decrease in functionally active ribonucleotide reductase. Thus, we supplemented the cell medium with Fe-citrate, and observed that the mtDNA copy number increases in *Gaa*-KO cells with iron repletion relative to WT levels (Figure 6C).

Having shown that mtDNA levels can be rescued in iron-deficient cells by Fe-citrate supplementation in the medium, we sought to test if the same principle was valid *in vivo*. Therefore, we tested if increasing the concentration of iron in the diet would avoid loss of mtDNA *in vivo*, as well as ameliorate the downstream consequences of iron-deficiency in *Gaa*-KO mice. The change in iron levels in the diet is most efficient when it occurs at weaning. Thus, upon separating the litters from the mothers (at post-natal day 21), we gave the weaned mice an iron-enriched diet (500mg Fe/Kg) or a control diet (179mg Fe/Kg). We followed the mice until they were two months of age, and measured mtDNA copy number in the cortex. While the GAA-KO mice treated with the control Na-citrate diet showed a decrease in mtDNA in cortex relative to the WT littermates, the *Gaa*-KO cohort fed iron-enriched chow had mtDNA copy number similar to the WT mice (Figure 6D). This result shows that the mitochondrial phenotype of young *Gaa*-KO mice can be rescued by increasing the levels of dietary iron.

### Dietary iron supplementation corrects inflammation in the *Gaa*-KO brain

The rescue of mtDNA levels in *Gaa*-KO brain by using an Fe-enriched diet encouraged us to test if other parameters due to iron deficiency would also be corrected, in particular inflammation. So, we analyzed the transcript levels of interferon-stimulated genes and those of their regulators in the cortex of 2-month old mice in Fe-enriched or control diet, and observed that in the iron-supplemented *Gaa*-KO mice the inflammatory response was ablated (Figure 6E). Furthermore, we observed that the protein levels of GFAP and of the inflammatory mediator TNFα were also decreased by the iron-supplementation in the *Gaa*-KO cortex (Figure 6F).

Next, we sought to characterize the molecules driving the inflammatory response. To achieve that, we employed a cytokine array, and used *Gaa*-KO and WT 2-month old cotical extracts, from mice that were either iron-supplemented or in the control diet. While we barely could observed TNFα, interferon gamma (IFNγ) as well as the cytokines CXCL9, CXCL10, CXCL11, IL-1α, IL-16 in wild type mice, these proteins were significantly increased in the cortices of *Gaa*-KO fed standard chow, but were normalized to the barely detectable WT levels in the iron-supplemented mice (Figure 6G, Supplementary Figure S5H). Other inflammatory molecules, including TREM-1, M-CSF, TIMP-1, C5, ICAM-1 and CXCL12 showed a similar profile (Supplementary Figure S5G). These results show that the pro-inflammatory phenotype observed in the brain of the *Gaa*-KO mice is due to the impaired iron homeostasis and can be corrected by increasing iron concentration in the diet.

To test whether the dietary iron supplementation had any effects on the behavior of the mice, we made a broad analysis of gait and motor parameters of the mice using the DigiGait platform (Rostosky and Milosevic, 2018). We observed that several parameters were affected in the *Gaa*-KO already at 2 months of age, particularly the stride length, which was decreased in the *Gaa*-KO mice (Figure 6H). This is an indication of gait disorder (Fernagut et al., 2002), suggesting that the *Gaa*-KO mice may have motor impairments. Notably, the *Gaa*-KO mice supplemented with iron showed a normalization of the stride length (Figure 6H). Taken together, normalization of iron levels by a Fe-enriched diet restores mtDNA stability and ablates inflammation at 2 months of age in a mouse model of lysosomal malfunction causing functional iron deficiency.

## DISCUSSION

This study shows that iron deficiency can cause necrotic cell death and inflammation in cultured cells as well as *in vivo*. Furthermore, we establish the role of (endo)lysosomal acidification as a key step in cellular iron homeostasis. Notably, the necrosis and inflammatory signaling triggered by (endo)lysosomal malfunction-induced iron deficiency can be ablated by supplementation of iron that bypasses the endo-lysosomal pathway.

We show here that Fe release from (endo)lysosomes to cytoplasm requires acidic pH. This step is key to maintaining appropriate cytoplasmic iron levels. When the (endo)lysosomes are not properly acidified, Fe^3+^ obtained from ferritinophagy, mitophagy or transferrin endocytosis cannot be reduced to Fe^2+^. This reduction is catalyzed by the ferroreductase STEAP3, which has an optimal acidic pH (Lambe et al., 2009). The inability to reduce Fe^3+^ to Fe^2+^ results in the accumulation of Fe^3+^ in the lysosomes, since only Fe^2+^ can be exported via the channels divalent metal transporter 1 (DMT1) or mucolipin-1 (MCOLN1) (Dong et al., 2008; Touret et al., 2003). In addition to the excess of iron in the lysosomes, there is lack of it in the cytoplasm. The proteins that sense cellular iron function in the cytoplasm implying that cytoplasmic Fe levels determine when the cellular iron deficiency response is activated. Low Fe levels result in the conversion of aconitase in IRP1 and stabilization of IRP2 (Rouault, 2015). Together, these proteins promote iron mobilization and uptake, while repressing its storage and export from the cell. This program is carried out by repressing the translation of ferritin heavy and light chains (to inhibit storage), and by stimulating the expression of transferrin receptor (TFRC), in order to increase the uptake of iron. This response keeps Fe within a tight range of concentrations, but we show here that it requires functional lysosomes. When lysosomes are defective, either by treatment with v-ATPase inhibitors or by genetic mutations that impair the acidification of a pool of the cellular lysosomes, the iron homeostatic cycle is also impaired because it cannot be released in sufficient amounts to the cytoplasm.

Iron is an essential microelement, and life relies on iron availability (Rouault, 2015). Accordingly, inhibition of lysosomal v-ATPase and the consequent iron deficiency result in cell death. It can be argued that this is due to toxicity of the inhibitors used, or to effects in other cellular organelles (the v-ATPase is also present at the Golgi and in synaptic vesicles) or processes. Nevertheless, the simple repletion of iron through an (endo)lysosome-independent pathway was sufficient to restore cell proliferation almost to control levels. Therefore, the main contributor to cell death caused by impaired (endo)lysosomal acidification seems to be iron deficiency. We determined that the cell death caused by iron deficiency is necrotic, which likely contributes to the inflammation observed in vivo in the brain of a mouse model of lysosomal malfunction (*Gaa*-KO). In addition, iron deficiency was sufficient to trigger inflammatory signaling in cultured primary cortical neurons. While we cannot exclude the possibility that necrotic death of cells in culture was sensed by astrocytes or glia that are present in the culture (primary neurons typically have a small number of co-cultured astrocytes and microglia), this result suggests that the inflammation trigger is cell-intrinsic.

Many cellular processes rely on iron, from DNA replication (the synthesis of dNTPs is Fe-dependent) to metabolism (mitochondrial respiratory chain relies on Fe-S clusters) (Rouault, 2015). Importantly, mitochondrial respiratory chain and oxidative phosphorylation subunits are encoded by two genomes, the nuclear DNA and the mitochondrial DNA (mtDNA). There is no dedicated *de novo* production of dNTPs for mtDNA synthesis, and mtDNA replication relies mostly on the cytoplasmic dNTP production (Copeland, 2012). A key step in dNTP production is catalyzed by the enzyme ribonucleotide reductase, a heterodimeric protein that has Fe in its active center (Puig et al., 2017). Decreased availability of dNTPs is a known cause of mtDNA instability, which in turn is an inflammatory trigger. We show here that iron deficiency perturbs mtDNA homeostasis, regardless of the cell type (fibroblasts, neurons) or the mechanism (genetic, pharmacologic) that leads to iron deficiency. The levels of mtDNA decrease sharply in iron deficiency. This effect is akin to what is observed in the mice lacking one allele of the mitochondrial transcription factor A (TFAM), a protein needed to replicate, transcribe and stabilize mtDNA (Ekstrand et al., 2004; West et al., 2015). In the *Tfam*^+/−^ mice, mtDNA instability triggers interferon-dependent signaling that culminates in potent inflammatory responses (West and Shadel, 2017). We show here that iron deficiency, either caused by iron chelation, genetic ablation of *Irp1* or *Irp2*, or genetic defects in the lysosomes, is associated both with mtDNA instability and inflammation, *in vitro* and, importantly, *in vivo*. Notably, supplying iron via an endo/lysosome-independent manner resolved the iron deficiency response, the loss of mtDNA and the inflammatory signaling.

Another process that depends on iron is the activity of the α-ketoglutarate-dependent superfamily of dioxygenases (Raimundo et al., 2011). These enzymes transfer oxygen atoms to different substrates, in a manner that is dependent on O_2_, Fe^2+^ and α-ketoglutarate. Absence of any of these factors, or accumulation of succinate (a product of these reactions) results in inhibition of the dioxygenases. The prolyl hydroxylases are part of the α-ketoglutarate-dependent superfamily, and catalyze the hydroxylation of the alphasubunit of hypoxia-inducible factor (HIF-1α). After hydroxylation, HIF-1α is recognized by the ubiquitin ligase VHL, and labeled for proteasomal degradation (Raimundo et al., 2011). HIF-1α is therefore constitutively expressed and readily degraded under normal conditions, and thus barely detectable (Raimundo et al., 2011). However, in the absence of VHL or when the function of prolyl hydroxylases is impaired, HIF-1α accumulates and triggers a transcriptional program known as (pseudo-)hypoxia response. Therefore, the activation of HIF signaling is a novel consequence of lysosomal dysfunction. Notably, HIF signaling can also be activated by mitochondrial malfunction, particularly of the citrate cycle enzymes fumarate hydratase or succinate dehydrogenase, which results in the accumulation of fumarate and succinate, and in the consequent inhibition of the prolyl hydroxylases. Therefore, HIF can be seen as a sensor of the mitochondrial-lysosomal axis and, reciprocally, the constitutive silencing of the (pseudo)hypoxia response requires both functional lysosomes and functional mitochondria. This aspect illustrates a specific mechanism of cooperation between different organelles, in this case, between mitochondria and lysosomes.

We and others have recently unveiled several signaling pathways by which deficient mitochondria result in lysosomal impairment, or vice-versa (Demers-Lamarche et al., 2016; Fernandez-Mosquera et al., 2017; Fernandez-Mosquera et al., 2019; Osellame et al., 2013; Yambire et al., 2019). This mutually destructive relationship between mitochondria and lysosomes is particularly detrimental in neurodegenerative diseases. For example, many of the genes that cause inherited Parkinson’s disease are associated either with mitochondria or lysosomes/autophagy, and therefore the primarily-damaged organelle is likely to promote the secondary impairment of the other. This manuscript presents a novel role of mitochondria-lysosome crosstalk in the regulation of central nervous system inflammation *in vivo*.

---

**Figure S1.**
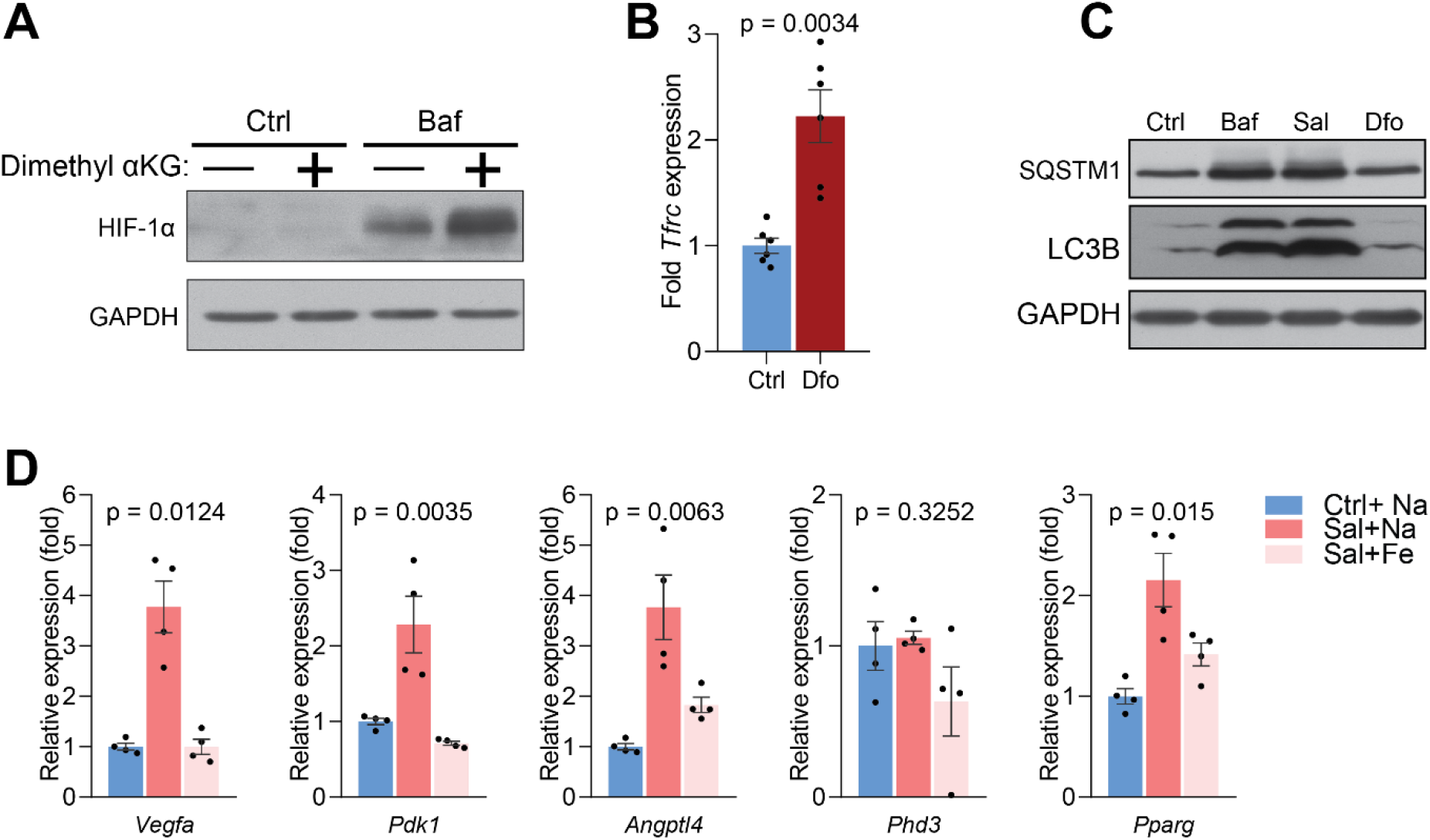
HIF-1a mediated response is triggered by lack of lysosomal iron efflux (related to Figure 1). (A)Whole cell Western blot of HIF-1a levels in control and Baf-treated cells supplemented with a cell permeant a-ketoglutarate. GAPDH is used as a loading control (n = 3). (B) Increased *Tfrc* mRNA levels in cells treated with lysosomal iron chelator, deferoxamine (Dfo) for 24 hours. Bar graphs represent mean±SEM for six independent experimental measures (shown as black dots). p value is unpaired two-tailed t-test with Welch’s correction (C) Immunoblots of SQSTM1 and LC3B in Baf- and Sal-treated cells (n = 6). Dfo is used as positive control for the independence of impaired lysosomal iron efflux on defective autophagy. GAPDH is used as loading control. (D) Expression level of HIF-1a target genes in Sal- and Sal-treated cells with iron supplementation. p values are estimated as Welch’s one-way ANOVA with all experimental groups compared to Sal-treated cells. Multiple test corrections were by the Dunnett’s method.

**Figure S2.**
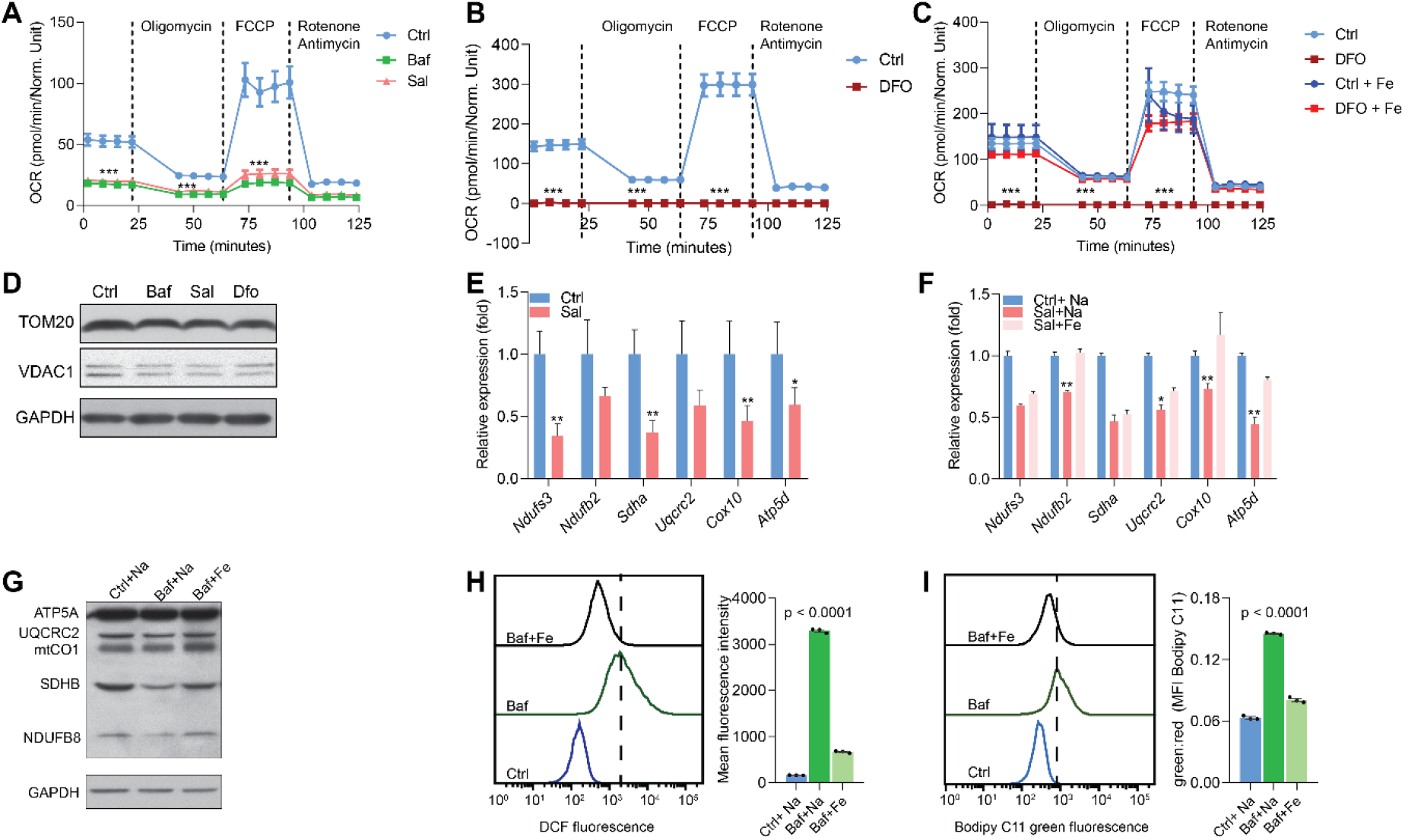
Lysosomal iron efflux is essential for mitochondrial biogenesis and function (related to Figure 2). (A-B) Mitochondrial OCR in Baf- and Sal-treated cells (A), and in Dfo-treated cells (B). Note the seemingly lack of mitochondrial respiration in Dfo-treated cells. Results represent mean±SEM, n =3. Each experimental replicate is calculated from the average of 12 technical replicates. ***p < 0.001, Welch’s one-way ANOVA with Dunnett’s correction for multiple comparisons (A) and unpaired two-tailed t-test with Welch’s correction (B). (C) Iron supplementation rescues the barely detected mitochondrial OCR in Dfo-treated cells. Experimental points represent mean±SEM, n =3. Each experimental replicate is calculated from the average of 8 technical replicates. ***p < 0.001, Welch’s one-way ANOVA with Dunnett’s correction for multiple comparisons with all experimental groups compared to Dfo-treated cells. (D) Immunoblots showing reduced TOM20 and VDAC1 protein levels in cells treated with Baf and Sal (n = 6). Dfo is used as positive control for iron deficiency and GAPDH is used as loading control. (E) Mitochondrial biogenesis is repressed in Sal-treated cells as assessed by the reduced transcript levels of nuclear-encoded mitochondrial genes following 24 hours of treatment with Saliphenylhalamide (Sal). Results are shown as bar graphs representing mean±SEM of six independent experimental replicates. *p < 0.05; **p <0.01; ***p < 0.001, unpaired two-tailed t-test with Welch’s correction. (F) Iron supplementation in Sal-treated cells, partially rescues the decreased expression of nuclear-encoded mitochondrial genes in cells treated with Sal alone. Bars depict mean±SEM, n = 8. *p < 0.05; **p <0.01; Welch’s one-way ANOVA with Dunnett’s correction for multiple comparisons. All conditions are compared to the Sal alone treatment group. (G) Whole cell immunoblots of ATP5A, UQCRC2, mtCO1, SDHB and NDUFB8 in Baf-treated cells and in iron supplemented-Baf treated cells (n = 3). Note the rescue of the decreased levels of mitochondrial proteins (NDUFB8, SDHB and mtCO1) following iron supplementation in Baf-treated cells. GAPDH is used as loading control. (H-I) Iron supplementation rescues Baf-induced increase in the levels of reactive of oxygen species (DCF) and in the levels of lipid peroxidation (Bodipy C11). Results are depicted representative histograms of three independent experimental replicates and quantifications are shown as mean±SEM. p values are determined by the Welch’s one-way ANOVA with Dunnett’s correction for multiple comparisons to Baf-treated cells.

**Figure S3.**
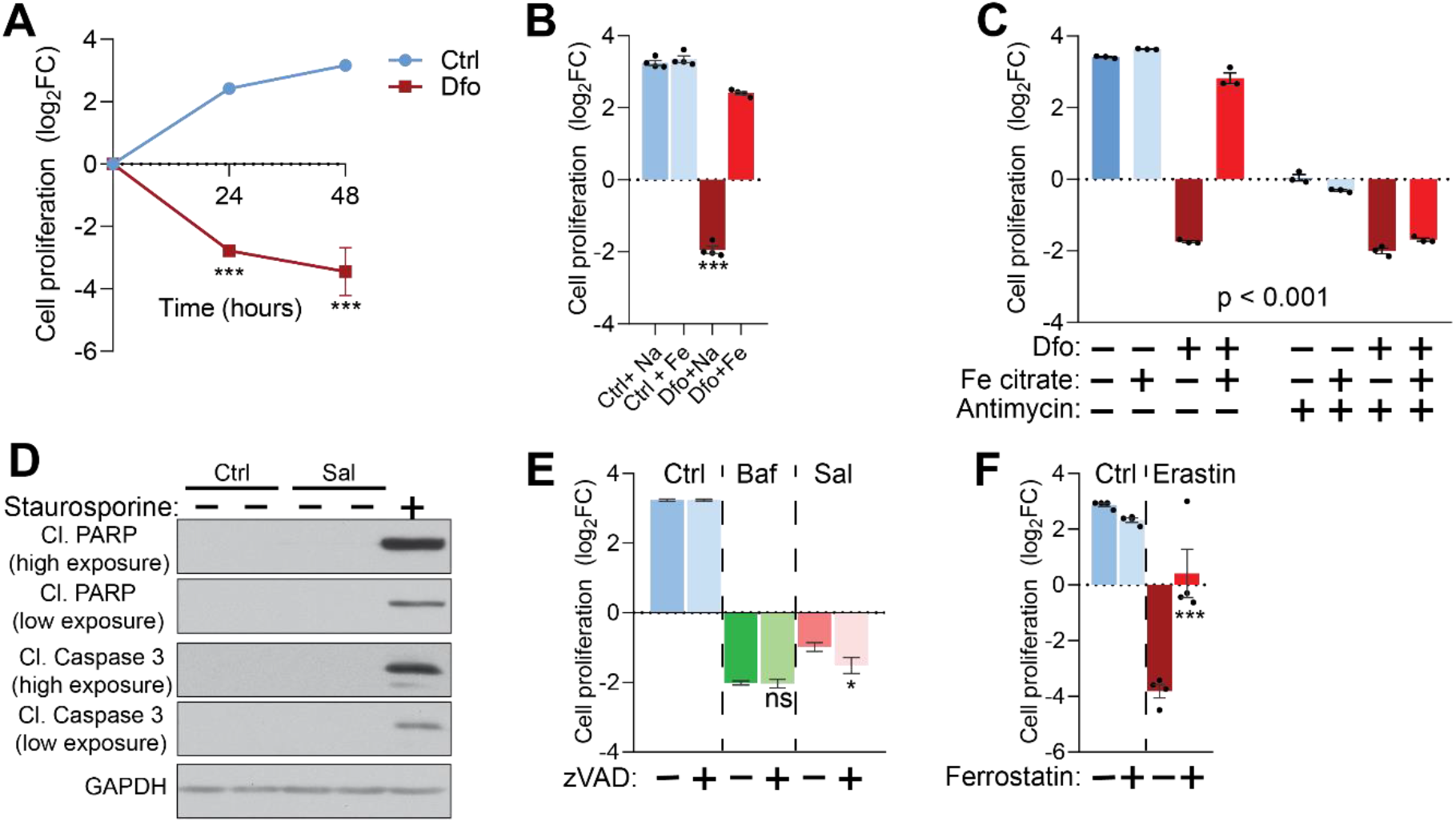
Iron deficiency-induced cell death is non-apoptotic (related to Figure 3). (A) Dfo-treated cells show time dependent increase in cell death of up to about sixteen-fold after 48 hours of treatment. Data points are mean±SEM, n = 4. ***p < 0.001, unpaired t-test with Welch’s correction (B) Iron supplementation rescues cell death following Dfo treatment. Bars and error bars represent mean±SEM of log2 fold change in cell number of experimental conditions. Black dots indicate individual experimental measures. *** p <0.001, Welch’s one-way ANOVA with Dunnett’s correction for multiple comparisons. Mean differences compared between Dfo alone treatments and the other experimental groups. (C) 20µM Antimycin treatment for 48 hours abrogates the rescue of cell death following iron supplementation in Dfo-treated cells. Results are summarized at mean±SEM, n = 3 (depicted as black dots). ***p < 0.001; Two-way ANOVA with Sidak correction of for multiple comparisons. (D) Caspase 3-independent cell death in Sal-treated cells shown by whole cell immunoblots of cleaved PARP and cleaved caspase 3 levels in Sal-treated cells (n = 3). Staurosporine (1µM treatment for 4 hours) is used as positive control for caspase 3-dependent cell death. GAPDH is used as loading control. (E) Treatment with 20µM zVAD, the pan-caspase inhibitor, for 48 hours results in no observable effects on cell death induced Baf or Sal treatment. Results show mean±SEM of four experimental replicates. *p < 0.05; ns p >0.05, Welch’s one-way ANOVA with Dunnett’s correction of multiple comparisons. Baf or Sal alone treated groups compared with all other treatment conditions (F) Ferrostatin treatment abrogates Erastin-induced cell death.Treatments were carried out with 5µM Ferrostatin and 10µm Erastin for 48 hours. Bar graphs represent mean±SEM, n = 4 (black dots). ***p < 0.001, Welch’s one-way ANOVA with Dunnett’s correction for multiple comparisons (all groups compared to cells treated with erastin alone).

**Figure S4.**
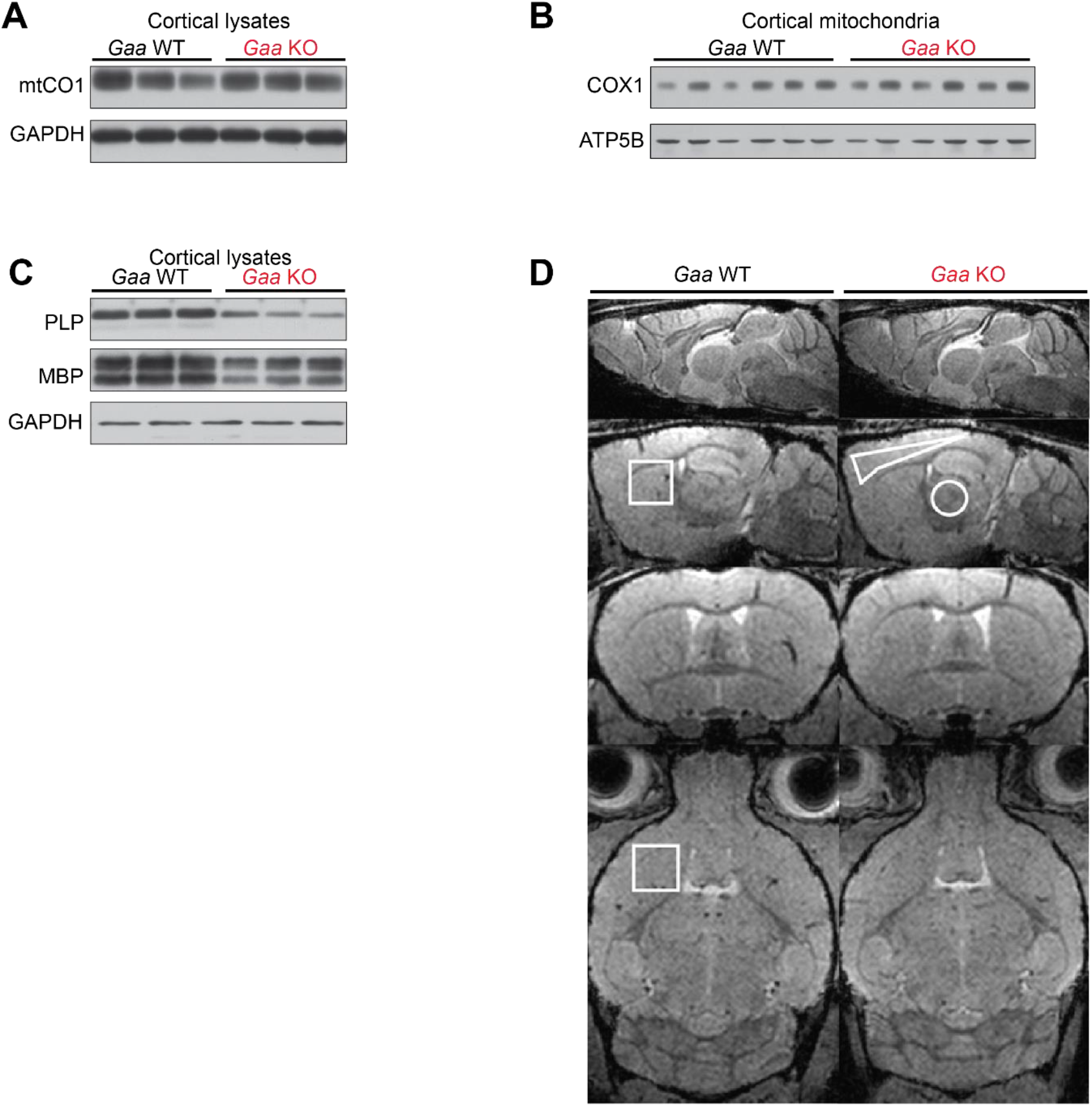
Steady state levels of mtCO1, PLP and MBP proteins (related to Figure 4). (A) Immunoblots of mtCO1 in whole cortical lysates of *Gaa* WT and KO mice at 6 months of age. GAPDH is used as loading control (n = 8). (B) Western blots of COX1 in mitochondria purified from *Gaa* WT and KO mouse cortices at 6 months of of age. ATP5B is used as loading control (n = 5). (C) PLP and MBP immunoblots from whole cortical lysates of 6-month old *Gaa*^−/−^ mice and their wild type littermates (n = 8). GAPDH is used as loading control. (D) MRI of the brain of a 6-month-old wild-type (left column) and a 6-month-old *Gaa*^−/−^ mouse (right column) in the mid-sagittal section (upper row), the sagittal section 2.4 mm from the midline (second row), a coronal section showing the anterior commissure (third row), and a horizontal section (bottom row). They show the volume-as well as the region-of-interest used for the evaluation of the striatum (squares), the cerebral cortex (polygon), and the thalamus (circle).

**Figure S5.**
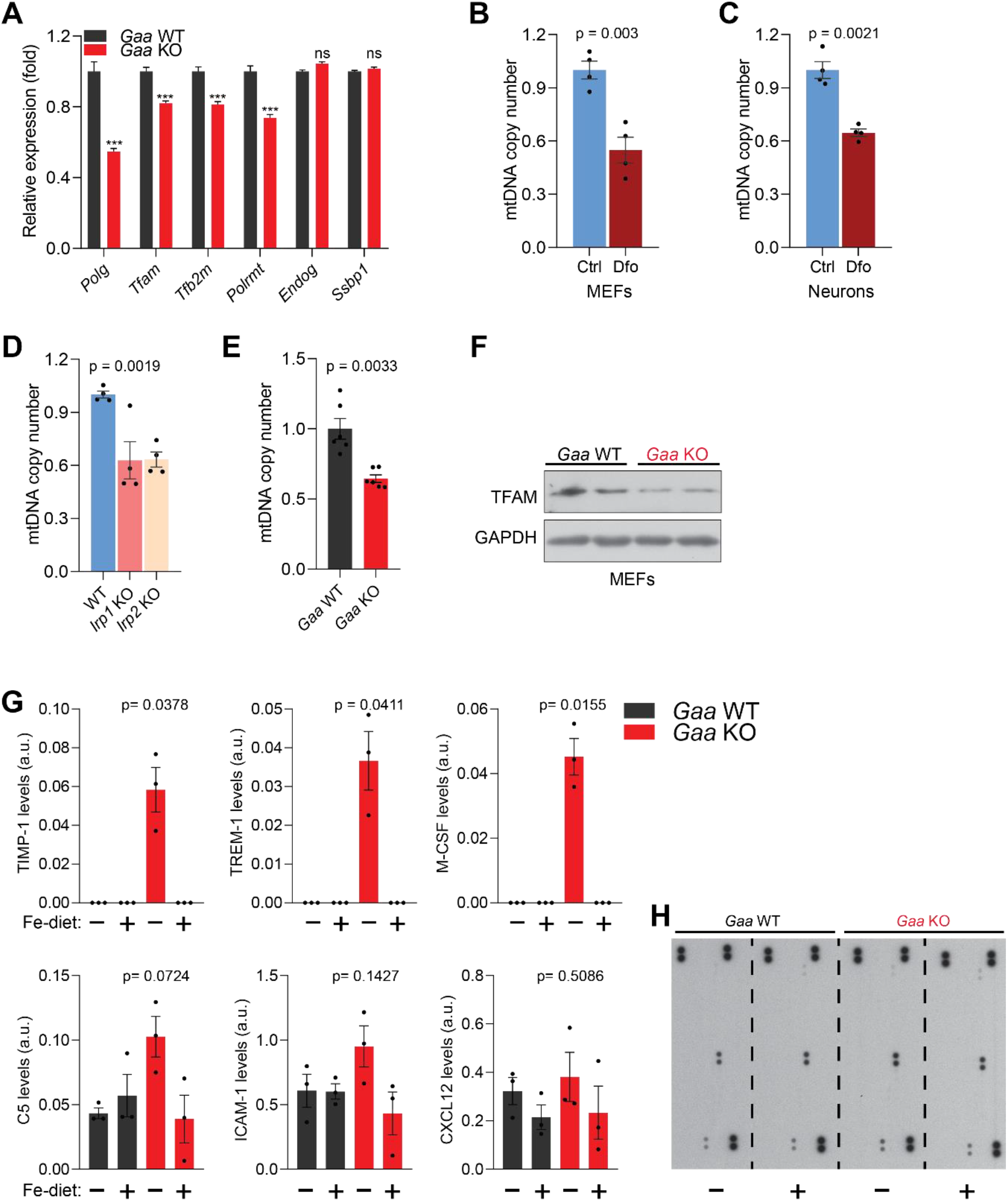
Iron deficiency causes impaired mtDNA homeostasis and induces inflammation in vivo (related to Figure 6) (A) Transcript levels of mtDNA maintenance and transcription genes in *Gaa*^−/−^ mice and their wild type littermates. Graphs represent mean±SEM, n = 8 mice per group. ***p < 0.001; ns p > 0.05, unpaired two-tailed t-test with Welch’s correction. (B-C) mtDNA copy number levels in Dfo-treated MEFs (B) and neurons (C). Results are summarized as mean±SEM of four independent experiments (depicted as black dots). p values are estimated as unpaired two-tailed t-test with Welch’s correction. (D) *Irp1*^−/−^ and *Irp2*^−/−^ MEFs show reduced mtDNA copy number levels. Bars represent mean±SEM, n = 4 (black dots). p value, Welch’s one-way ANOVA with Dunnett’s correction for multiple comparisons made to WT group. (E) mtDNA copy number decline in *Gaa*^−/−^ MEFs relative to WT. Results are shown as mean±SEM, n = 6 (black dots). p value is the unpaired two-tailed t-test with Welch’s correction. (F) Whole cell immunoblots of TFAM in *Gaa*^−/−^ MEFs and their WT controls (n = 4). GAPDH is used as a loading control. (G) Iron-enriched diet restores increased levels of cytokine and chemokines to control levels in the cortices of 2-month old *Gaa*^−/−^ mice. Cytokines are increased in *Gaa*^−/−^ mice fed standard chow, and are barely detectable in *Gaa*^−/−^ mice fed iron-enriched diet or their wild type littermate controls. Levels of cytokines and chemokines are shown as mean±SEM (n = 3 mice per group, black dots) and p values represent Brown-Forsythe’s one-way ANOVA tests estimated as differences between *Gaa*^−/−^ mice on standard chow and all other experimental groups (H) Representative image of cytokine array blots (n = 3animals per group) quantified and presented as graphs in Figure 6G and Figure S5G.

## METHODS

### EXPERIMENTAL MODEL AND SUBJECT DETAILS

#### Mice

The *Gaa*^−/−^ mice (B6:GaatmRbn/J) used in this study were purchased from the stock of The Jackson Laboratory, bred and housed under standard pathogen-free conditions at the animal facility of the European Neuroscience Institute with access to food and water *ad libitum*. All animal experiments were carried out in accordance with the European guidelines for animal welfare and were approved by the Lower Saxony Landesamt fur Verbraucherschutz and Lebensmittelsicherheit (LAVES) registration number 15-883. Except for iron-enriched chow experiments in which mixed sex cohorts were used at 2 months of age, all other experiments were performed on only male cohorts at 0, 2, 6 and 12 months of age. Regular chow contains 179mg Fe/Kg, and the iron-enriched chow contains 500mg Fe/Kg. The iron-enriched and control diet were purchased from Brogaarden (Denmark).

#### Cell lines and primary cultures

Murine embryonic fibroblasts were prepared in-house from *Gaa*^−/−^ mice and their wildtype littermate controls. HeLa (female ATCC CCL-2) were obtained from ATCC. MEFs from wild type, *Irp1*^−/−^ and *Irp2*^−/−^ mice were obtained from the Technion-Israel Institute of Technology (Esther Meyron Holtz). All these lines were cultured in Dulbecco’s modified Eagle’s medium with high glucose (Gibco) supplemented with 10% fetal bovine serum (FBS) and 1% Penicillin/Streptomycin at 37°C and 5% CO_2_, in a humidified incubator, unless otherwise stated. Wild type C57BL/6J P0 pups were used to prepare primary cortical cultures as previously described (Ferguson et al., 2007). Briefly, mouse cortices were dissected under sterile conditions and digested in freshly prepared, warm enzyme solution (1.5mM cysteine, 20U/mL papain, 0.75mM EDTA, 1.5mM CaCl2 and 10ul/mL DNase) in HBSS medium for 30 minutes at 37°C on a shaker. The enzyme solution was replaced with plating medium (Neurobasal medium supplemented with 5% FBS, 1% glutamax, 2% B27 supplement and 0.5% Penicillin/Streptomycin) for 5 minutes. Neuronal tissue was washed in HBSS medium and cultured in plating medium for 12 hours at 37°C and 5% CO_2_, in a humidified incubator. The plating medium was replaced with neuronal medium (as plating medium without 5% FBS) and neurons were cultured under standard growth conditions (37°C and 5% CO_2_) in a humidified incubator. All neuronal experiments were carried out with neurons at DIV 14.

### METHOD DETAILS

#### Behavior experiments

Mouse behavior experiments were performed as detailed in Rostosky and Milosevic (2018). In brief, ventral plane videography for gait analysis was performed on *Gaa*^−/−^ mice and their wild type littermate controls fed standard or iron-enriched chow for 2 months using Digigait instrumentation at a speed of 40cms^−1^. For each group 13-19 male mice were used for behavior experiments. Results were analyzed with software from Mouse Specifics according to the manufacturer’s protocol and differences were determined by the Welch’s one-way ANOVA test with Dunnett’s correction for multiple comparison to *Gaa*^−/−^ mice fed standard diet.

#### Cell culture drug treatments

All experiments involving drug treatments or iron supplementation in cells were carried out under the standard growth conditions stated above (see “Cell lines and primary cultures”) unless otherwise stated. Briefly, following seeding and overnight incubation, cells were washed with warm PBS and treated for 24 hours with growth medium containing final concentrations of 0.5µM bafilomycin, 0.5µM saliphenylhalamide or 300µM deferroxamine. Experiments involving iron supplementation were carried out for 48hours with a final concentration 150µM Fe-citrate (or Na-citrate as control) in growth medium.

#### Cell proliferation assay

Cell proliferation was determined using the Cell Titer Glo Luminiscence cell viability assay (Promega)following manufacturer’s instructions. Briefly, 8000 cells per well were cultured in 200µL of DMEM high glucose medium supplemented with 10% FBS and 1% Penicillin/Streptomycin as at least technical triplicates for each treatment in a 96-well plate. Following overnight culture, the initial number of cells prior to treatment was determined from five untreated wells. Cells were then treated with the indicated drugs as described in each experimental condition for 24 hours or 48 hours in 100µL of medium. Prior to cell number measurements, plates and reagents were equilibrated to room temperature. 100µL of Cell Titer Glo reagent (Promega) was added to each well, mixed and the luminescence after 10 minutes of incubation was read in a microplate reader (Synergy H1, Biotek). Cell proliferation rates determined from a standard calibration curve and reported as log_2_ fold change of cell numbers relative to that at the initial time point prior to treatment.

#### Mouse brain sample preparation

For all biochemical and q-RT-PCR experiments, mouse tissues were harvested, snap frozen in liquid nitrogen and stored at −80°C. Briefly, mice were anesthesized with isofluorane and euthanized by cervical dislocation. Mouse brain was dissected into cortices, hippocampi and rest of brain, rinsed in PBS, transferred into tubes and snap frozen in liquid nitrogen. These samples were frozen at −80°C until processed for each experiment accordingly. For Immunohistochemical assays, mice were anethesized by peritoneal injection with 40mg/mL chloral hydrate in PBS and perfused with 100U/mL heparin followed by cold 4% *para*-formaldehyde (PFA) in PBS. Harvested brains were post-fixed overnight in 4% PFA at 4°C, slowly dehydrated in 12-18% gradients of sucrose solutions and embedded in OCT (Tissue Tek) in a cryomold, frozen in isopentane set in liquid nitrogen and stored at −80°C.

#### Fluorescence microscopy and image analysis

To determine cytoplasmic and mitochondrial labile iron levels by microscopy, cells were seeded in poly-L-lysine coated coverslips in 12-well plates, treated with bafilomycin for 24 hours and loaded with FerroOrange and Mito-FerroGreen respectively. Live cell imaging of cells was performed with a Nikon/Perkin Elmer Ultra VIEW VoX system (Perkin Elmer). Mitochondrial morphology of bafilomycin-treated cells was assessed as described (Fonseca et al., 2019). Briefly, cells were electroporated (Amaxa Nucleofector) with pAcGFP1-Mito and seeded into poly-L-lysine coated coverslips in 12-well plates, treated for 24 hours with 0.5uM Bafilomycin and fixed with 3.7% PFA for 15 minutes at room temperature. About 0.2µm thick Z-stacks were obtained with the 63x objective of the LSM800 Airyscan setup (Zeiss). Frozen brain tissue as prepared before see “Mouse brain sample preparation” above were used for Immunohistochemical assays to determine GFAP levels. In brief, 10µM thick sagittal brain sections were cut with a cryostat, post-fixed and permeabilized with 0.1% Triton X-100. Immunofluorescence of the sections was performed as described (Milosevic et al., 2011). Image analysis was carried out as detailed in Fonseca et al. (2019) using ImageJ software version 1.51j8.

#### Isolation of cortical mitochondria

Mouse cortices were harvested and mitochondria were purified from the tissues as described in (Lazarou et al., 2007). In brief, tissues were Dounce homogenized on ice in homogenization buffer (300mM Trehalose, 10mM KCl, 10mM HEPES pH 7.4 supplemented freshly with 0.2% BSA and protease/phosphatase inhibitor cocktail), centrifuged twice at 700xg for 10 minutes at 4°C to pellet tissue debris and nuclear fractions. The supernatant was centrifuged at 11000xg for 15 minutes at 4°C to separate the mitochondria.

#### Blue native (BN) –PAGE and complex IV activity staining

Isolated mitochondria were solubilized in buffer containing 1% digitonin and protein complexes were separated by BN-PAGE as previously described (Witting and Shägger, 2005).. For native complex IV activity staining, gels were equilibrated in 50mM Potassium phosphate buffer (pH 7.4) and stained at 30°C with complex IV activity staining buffer (50mM KPi pH 7.4, 0.5mg/mL diaminobenzidine (DAB) and 1mg/mL reduced cytochrome *c*) until the brown oxidized DAB stains were visible. The intensity of DAB stain (complex IV activity) was quantified with Imagequant TL (GE Healthcare Life Sciences). For native complex V activity staining, gels were equilibrated in 35mM Tris, 220 mM Glycine (pH 8.3) and stained at 30°C in assay buffer (35 mM Tris, 220 mM glycine, 14 mM MgSO_4_, 0.2% Pb(NO_3_)_2_, 8 mm ATP (pH 8.3)).

#### Immunoblotting

Whole cell extracts of cultured cells were lysed in 1.5% n-dodesylmaltoside (Roth) in PBS supplemented with protease and phosphatase inhibitor cocktail (Thermo Fisher Scientific) as described (Fernandez-Mosquera et al., 2019). For brain (cortical) lysates, snap frozen and ground tissues were lysed in RIPA buffer (50mM Tris pH 8.0, 150mM NaCl, 1% Triton X, 0.5% sodium deoxycholate and 0.1% SDS) freshly supplemented with protease and phosphatase inhibitor cocktail. Protein concentrations of whole cell extracts and tissue lysates were determined using a Bradford Protein assay (Bio-Rad). 50μg of sample proteins per well were subjected to SDS-PAGE and transferred to polyvinylidene fluoride (PVDF) membranes (Amersham, Life Technologies). Membranes were blocked in 5% Milk in TBS tween and probed with the following primary antibodies: HIF-1 alpha (Novus), GAPDH (Sigma-Aldrich),SQSTM1 (Abcam, ab110252), HPRT (Abcam, ab10479), (LC3B (Cell signaling, 3868), mtTFA (Abcam, ab138351), Total rodent OXPHOS cocktail (Abcam, ab110413), TNFα (Abcam), cleaved caspase 3 (Cell signaling), cleaved PARP (Cell signaling), GFAP (Abcam), PLP and MBP (Kind gift of K-A Nave, MPI-EM), FTH1 (Cell signaling), Ferritin light chain (Abcam), TOM20 (Proteintech), VDAC1 (Abcam), VHL (Santa Cruz) and self-made rabbit polyclonal antibodies to COX1 and ATP5B. HRP-conjugated secondary antibodies against mouse and rabbit IgGs (The Jackson Laboratory) were used. Band densitometric quantifications were determined using Image J software version 1.51j8.

#### Mitochondrial respiration measurements

Mitochondrial oxygen consumption rate (OCR) was measured in cells using the XF96 Extracellular Flux analyzer (Seahorse Bioscience) as described (Yambire et al., 2019). Briefly, cells were seeded at 20000 cells per well in XF96 cell culture multi-well plates and cultured overnight under standard conditions stated previously (see “Cell lines and primary cultures” above). Cells were then treated with the indicated drugs for 24 hours while XF96 cartridges were incubated overnight in XF calibrant at 37°C in a non-CO2 incubator. Prior to OCR measurements, the growth medium containing the indicated treatments of cells was exchanged with XF medium containing 1µg/mL Hoechst dye, stained for 10 minutes to determine cell numbers, and subsequently incubated at 37°C in a non-CO2 incubator for 1hour. For OCR profiles, 1uM Oligomycin, 2uM FCCP and 1uM mix of Rotenone and Antimycin loaded into corresponding microwells in the XF96cartridge plate. Following equilibration of sensor cartridges, XF96 cell culture plate was loaded into the XF96 Extracellular Flux analyzer at 37°C and OCR was measured after cycles of mixing and acquiring data (basal) or inhibitor injection, mixing and data acquisition. Results were analyzed by the WAVE desktop software (Agilent) and normalized to the number of cells determined prior to assay.

#### Quantitative RT-PCR

RNA from cells was extracted using the Crystal RNA mini Kit (Biolab,). For tissue RNA extraction, the TriReagent (Sigma, T9424) was used according to supplier instructions. RNA concentration and quality were determined using Nanodrop (PeqLab) and cDNA was synthesized with the iScript cDNA synthesis kit (Bio-Rad) following manufacturer’s protocol. Each 9μl reaction for q-RT-PCR was made of 4μl diluted cDNA, 0.25μl of each primer (from 25μM stock) and 4.5μl of Luna Universal Master Mix (New England Biolabs). The q-RT-PCR reactions were run on the QuantStudio 6 Flex Real-Time PCR system (Applied Biosystems). qPCR results were analyzed using the ΔΔCt method relative to the mean of housekeeping genes (*Rps12*, *Hprt* and *Gapdh*). Relative values for mtDNA and nDNA genes were used to generate mtDNA copy number levels per nuclear genome. Each biological data point represents the average of at least technical triplicates.

#### mtDNA copy number analysis

To measure relative mtDNA copy number levels, total DNA was isolated from cultured cells or tissues as described previously in Cotney et al. (2007). Briefly, cells or tissues were lysed in 500µL DNA extraction buffer (50mM Tris-HCl pH 8.5, 0.25% SDS, 1mM EDTA pH 8.0, 5mM DTT) by boiling for 10 minutes. Samples were cooled to room temperature and incubated with 100µg RNAse A for 3 hours at 37°C followed by incubation with 100µg of Proteinase K at 55°C overnight. After Proteinase K digestion, samples were boiled at 95°C, allowed to cool to room temperature and the DNA concentration determined using Nanodrop (Peqlab). Diluted DNA samples were used along with mtDNA and nuclear DNA primers for qPCR as described above to determine relative mtDNA levels per genome.

#### Flow cytometry

Measurements of mitochondrial superoxide levels, lipid peroxidation and cellular reactive oxygen species (ROS) levels were determined using MitoSox^TM^ Red Superoxide indicator, Bodipy^581/591^ C11 and H_2_DCFDA (Thermo Fisher Scientific) respectively. Briefly, following treatments of cultured cells with indicated drugs for 24 hours, cells were washed in prewarmed PBS and loaded with either 1µM MitoSox, 1µM Bodipy^581/591^ C11 or 10µM H_2_DCFDA during 30 minutes in the dark. Control cells were treated with H_2_O_2_ (5mM) or Antimycin (20µM) for 20minutes as positive controls for increased levels of mitochondrial superoxides, lipid peroxidation or ROS. The mean fluorescent intensities corresponding to steady state levels of mitochondrial superoxides (Mitosox), lipid peroxidation (Bodipy^581/591^ C11) or ROS levels (H_2_DCFDA) were determined by flow cytometry (FACs Canto^TM^ II, BD Biosciences). Data were collected from 20,000 cells and results were analyzed by FlowJo vX.0.7 (Tree Star Inc.).

#### Measurement of cellular iron levels

Determination of cellular total, ferric and ferrous iron levels were carried out using the Iron assay Kit (Abcam) according to manufacturer’s instructions. Cells were washed in prewarmed PBS, scraped into ice-cold PBS and pelleted at 1000xg for 5minutes. Cell pellets were homogenized in assay buffer on ice using a Dounce homogenizer, centrifuged at 16000xg for 10minutes and the supernatant collected for the iron assay. 25µL of samples were made up to 100µL in a 96-well plate with assay buffer, and incubated for 30 minutes at 37°C with 5µL iron reducer (for total iron) or assay buffer (for ferrous iron) along with standards. 100µL of iron probe was added to each reaction, mixed and incubated for a further 1 hour at 37°C in the dark. The absorbance at 593nm, which corresponds to the iron levels, was determined using a microplate reader (Synergy H1, Biotek). Iron concentrations were calculated from the standard curve and normalized to the amount of protein for each sample, determined by the Bradford protein assay. The average of technical triplicates was used for each biological sample.

#### Magnetic resonance spectroscopy (MRS) and magnetic resonance imaging (MRI)

Mice underwent MRS, T_1_, T_2_, and MTR measurements of the cerebral cortex, striatum, and thalamus at 37ºC. After induction of anaesthesia by intraperitoneal injection of ketamine (100 mg/kg b.w.) and xylazine (10 mg/kg b.w.), animals were intubated with a purpose-built polyethylene endotracheal tube (0.58 mm inner diameter, 0.96 mm outer diameter) and artificially ventilated using an animal respirator (TSE, Bad Homberg, Germany) with a respiratory rate of 25 breaths per minute and an estimated tidal volume of 0.35 ml under 0.5% isoflurane as previously described (Watanabe et al., 2016). The animals were then placed in a prone position on a purpose-built palate holder equipped with an adjustable nose cone. Respiratory movement of the abdomen as well as rectal temperature were monitored by a unit supplied by the manufacturer (Bruker Biospin MRI GmbH, Ettlingen, Germany). At 9.4 T (Bruker Biospin MRI GmbH, Ettlingen, Germany), localized proton MRS (STEAM, TR/TE/TM = 6000/10/10 ms) was performed with the use of a birdcage resonator (inner diameter 70 mm) and a saddle-shaped quadrature surface coil (both Bruker Biospin MRI GmbH, Ettlingen, Germany) on anesthetized mice as previously described (Watanabe et al., 2016). Shimming of the B_0_ field was carried out by FASTMAP (Gruetter, 1993). A (20 mm)^3^ volume-of-interest was centered on the right striatum (squares in Supplementary Figure S4D). Water saturation was accomplished by means of three Gaussian-shaped CHESS radiofrequency (RF) pulses (90°-90°-180°), each of which with a duration of 7.83 ms and a bandwidth of 350 Hz. Overall duration of the CHESS module was 147 ms. Each CHESS pulse was followed by an associated spoiler gradient and a 37 ms outer volume saturation module covering a range of 3 mm around the volume-of-interest without gap. Metabolite quantification involved spectral evaluation by LCModel (Provencher, 1993) and calibration with brain water concentration of 79% (Duarte et al., 2014), for which the unsuppressed water proton signal served as internal reference. Metabolites with Cramer-Rao lower bounds above 20% were excluded from further analysis. Relaxation times and magnetization transfer ratios are determined in selected regions of the brain. T_2_ relaxation times of water protons were determined by a multi-echo spin-echo MRI (TR/TE=2500/10–123 ms). T_1_ relaxation times were determined with the use of a spin-echo saturation recovery sequence and 7 TR values from 0.15 to 6 s. For measurements of the magnetization transfer ratio, an off-resonance Gaussian-shaped RF pulse with a frequency offset of −3.6 kHz, duration of 8.5 ms, and a flip angle of 128° was incorporated into a spin density-weighted gradient-echo MRI sequence (RF-spoiled 3D FLASH, TR/TE = 19/4.2 ms, flip angle 5°, measuring time 97 s) at a resolution of 200×200×400 µm^3^. The duration and power of the off-resonance pulse was optimized to observe the transfer of saturation from non-water protons to water protons in brain of mice *in vivo* (Natt et al., 2003; Watanabe et al., 2012). For the evaluation of MRI signal intensities, square-shaped regions of interest with were selected in a standardized manner in the cerebral cortex (98 pixels), the striatum (100 pixels), and the thalamus (61 pixels) (Supplementary Figure S4D). The analysis followed a strategy previously developed for intraindividual comparisons of MR images obtained after manganese administration (Watanabe et al., 2004). Volumetric assessments were obtained by analyzing proton-density-weighted images (RF-spoiled 3D FLASH, TR/TE = 22/7.7 ms, flip angle 10°, fat suppression 90°, 117 µm isotropic resolution, measuring time 12 min, Supplementary Figure S4D) using software provided by the manufacturer (Paravision 5.0®, Bruker Biospin, Ettlingen, Germany). After manually outlining the whole brain and the ventricular spaces in individual sections, respective areas were calculated (in mm^2^), summed and multiplied by the section thickness. Significant differences between two groups of data were determined by Mann-Whitney’s U-test.

#### Cytokine measurements

Brain tissue cytokine levels were measured using the Mouse cytokine array Panel A (R&D Systems) following protocols described by the manufacturer. Briefly cortices from *Gaa*^−/−^ mice mice and their wild type littermate controls, fed iron-enriched or standard chow for 2 months were harvested, snap frozen in liquid nitrogen and ground using a tissue grinder. Tissues were lysed in RIPA buffer and spun at 10000xg for 5 minutes. The supernatants were collected and their protein concentrations determined by the BCA protein assay (Thermo Fisher Scientific). Nitrocellulose membranes containing capture antibodies were blocked in array buffer 6 for 1 hour. Samples were prepared by diluting 1mg of tissue lysates in 500µL array buffer 4 containing 15µL Detection antibody cocktail and made up to 1.5mL with array buffer 6. Block buffer on membranes was replaced with prepared samples. Following overnight incubation at 40C, membranes were washed three times in wash buffer for 10 minutes each, incubated for 30 minutes with Strepatavidin-HRP on a rocking platform, washed three more times and incubated for 1 minute with the Chemi Reagent mix. Membranes were developed onto X-ray films by a chemiluminiscence developer (AGFA) and the pixel density (using ImageJ) of each spot representing the level of the corresponding cytokine determined and normalized to that of the average of reference spots.

### QUANTIFICATION AND STATISTICAL ANALYSIS

#### Statistical analysis

All statistical analyses were carried out using Prism version 8.0 software. Unless otherwise stated in figure legends, to compare means between two groups, a two-tailed unpaired *t*-test with Welch’s correction was used for normally distributed data. Non-parametric, two-group means were compared by a two-tailed Mann-Whitney test. Welch’s one-way ANOVA, followed by Dunnett’s test for multiple comparisons was performed for multi-group (at least three) comparisons. For data on cytokine array experiments, the Brown-Forsythe’s one-way ANOVA test was used given the identical values in at least one of the columns. Measures were summarized as graphs displaying mean±S.E.M, of at least three independent biological replicates. Means of control samples are typically centered at one (or 100%) to ensure easier comparisons unless otherwise noted. Estimated p values are either stated as the actual values or denoted by * p<0.05; ** p<0.01; ***p<0.001. Differences were only considered as statistically significant when the p value was less than 0.05.

#### Data and software availability

The microarray and RNA sequencing data analysis was performed using Partek Software Suite. Ingenuity Pathway Analysis was used for assessment of transcription factor activity, as described in Yambire et al (2019). The RNAseq data generated in this study from the *Gaa*-KO and WT cortices is deposited in Gene Expression Omnibus.

## ACKNOWLEDGEMENTS

This work was supported by a Starting Grant from the European Research Council (337327, MitoPexLyso), to NR. This project stems from results obtained under the support of a research grant from the Acid Maltase Deficiency Association (to NR). IM lab was supported by a Emmy Noether award from the Deutsche Forschungsgemeinschaft (DFG). NR and IM were supported by the Collaborative Research Center SFB1190-P02 by DFG. The authors thank the technical support of Dirk Schwitters, and thank Adrian M. Raimundo for discussions on the project. We thank Prof. Dörthe Katschinski for the primers to test HIF targets.

## AUTHOR CONTRIBUTIONS

KFY suggested and performed most experiments, analyzed and interpreted data and prepared figures. TW performed the MRI experiments and analyzed the respective data together with JF. CR performed the behavioral experiments and assisted with the brain section immunostainings. IM, DP, ASG and PW suggested and performed experiments, analyzed and interpreted data. EMH shared key experimental reagents, helped define methodologies, analyzed and interpreted data. NR designed the study, performed experiments, analyzed and interpreted data and wrote the manuscript, which all authors then commented and edited.

## SUPPLEMENTARY FILES

Table S1. Annotation of disease or functions (Ingenuity Pathway Analysis) for the category of cell death and survival, for the differentially-expressed gene list obtained from dataset GSE47836.

Table S2. Annotation of disease or functions (Ingenuity Pathway Analysis) for the category of cell death and survival, for the differentially-expressed gene list obtained from dataset GSE60570.

Table S3. Annotation of disease or functions (Ingenuity Pathway Analysis) for the category of cell death and survival, for the differentially-expressed gene list obtained from dataset GSE16870.

Table S4. Concentration and concentration ratios of metabolites determined by *in vivo* MRS.

Table S5. List of qPCR primers for mouse genes.

