## Supplemental Table S4 for "Impaired lysosomal acidification triggers iron deficiency, necrotic cell death and inflammation *in vivo*"

Table S4. Concentration and concentration ratios of major metabolites determined by MRS (related to Figure 4)

|  |  | **6 months** | |  | **12 months** | |
| --- | --- | --- | --- | --- | --- | --- |
| Background |  | Wild type | *Gaa^-/-^* |  | Wild type | *Gaa^-/-^* |
| *n* |  | *n* = 8 | *n* = 8 |  | *n* = 5 | *n* = 5 |
| **tCr** |  | 11.5 ± 0.5 | 10.7 ± 0.3** |  | 13.3 ± 0.5 | 12.0 ± 2.4 |
| **GPC+PCh** |  | 3.8 ± 0.3 | 3.3 ± 0.2* |  | 4.5 ± 0.4 | 3.8 ± 0.6 |
| **GPC** |  | 1.9 ± 0.5 | 1.4 ± 0.7 |  | 1.7 ± 0.3 | 1.7 ± 0.2 |
| **PCh** |  | 1.9 ± 0.7 | 1.9 ± 0.6 |  | 2.7 ± 0.3 | 2.1 ± 0.4* |
| **NAA** |  | 8.7 ± 0.7 | 8.3 ± 0.7 |  | 10.0 ± 0.3 | 8.7 ± 1.6 |
| **Glu** |  | 11.7 ± 1.3 | 11.1 ± 0.7 |  | 12.2 ± 0.6 | 11.1 ± 3.0 |
| **Ins** |  | 3.4 ± 0.6 | 3.5 ± 1.0 |  | 2.9 ± 0.9 | 2.1 ± 1.1 |
| **Tau** |  | 14.7 ± 1.1 | 13.9 ± 1.9 |  | 15.2 ± 0.9 | 13.6 ± 1.9 |
| **GPC+PCh/tCr** |  | 0.33 ± 0.04 | 0.31 ± 0.02 |  | 0.34 ± 0.03 | 0.32 ± 0.03 |
| **GPC/tCr** |  | 0.16 ± 0.03 | 0.13 ± 0.06 |  | 0.13 ± 0.02 | 0.14 ± 0.01 |
| **PCh/tCr** |  | 0.17 ± 0.06 | 0.18 ± 0.06 |  | 0.21 ± 0.02 | 0.17 ± 0.03* |
| **NAA/tCr** |  | 0.75 ± 0.07 | 0.77 ± 0.07 |  | 0.75 ± 0.03 | 0.72 ± 0.06 |
| **Glu/tCr** |  | 1.0 ± 0.1 | 1.0 ± 0.1 |  | 0.92 ± 0.05 | 0.93 ± 0.2 |
| **Ins/tCr** |  | 0.29 ± 0.06 | 0.33 ± 0.09 |  | 0.22 ± 0.07 | 0.17 ± 0.08 |
| **Tau/tCr** |  | 1.3 ± 0.08 | 1.3 ± 0.2 |  | 1.1 ± 0.07 | 1.1 ± 0.2 |
| ***T_1_* (s)** |  | 1.62 ± 0.05 | 1.64 ± 0.07 |  | 1.62 ± 0.03 | 1.58 ± 0.06 |
| ***T_2_* (ms)** |  | 39.7 ± 1.0 | 37.3 ± 0.8** |  | 38.5 ± 0.6 | 36.7 ± 0.7* |
| ***MTR (%)*** |  | 9.9 ± 0.7 | 9.0 ± 0.8* |  | 9.8 ± 0.6 | 10.0 ± 0.9 |

Glu = glutamate, GPC+PCh = Choline-containing compounds, Ins = myo-inositol, MTR = magnetization transfer ratio, NAA = *N*-acetylaspartate, Tau = taurine, tCr = creatine + phosphocreatine. *, ** = p < .05, p < .01 vs. Wild Type in Mann-Whitney´s U-test.
