## Supplemental Table S5 for "Impaired lysosomal acidification triggers iron deficiency, necrotic cell death and inflammation *in vivo*"

Table S5 List of mouse qPCR primer sequences (related to Star methods)

| Oligonucleotide | SOURCE | IDENTIFIER |
| --- | --- | --- |
| HPRT qPCR  5’– CCTCCTCAGACCGCTTTTT – 3’  3’– AACCTGGTTCATCATCGCTAA – 5’ | This paper | N/A |
| RPS12 qPCR  5’– GAAGCTGCCAAGGCCTTAGA – 3’  3’– AACTGCAACCAACCACCTTC – 5’ | This paper | N/A |
| GAPDH qPCR  5’– TGTGTCCGTCGTTCTGA – 3’  3’– CCTGCTTCACCACCTTCTTGA – 5’ | This paper | N/A |
| TFRC qPCR  5’– GTTTCTGCCAGCCCCTTATTAT – 3’  3’– GCAAGGAAAGGATATGCAGCA – 5’ | This paper | N/A |
| FTH1 qPCR  5’– CAAGTGCGCCAGAACTACCA – 3’  3’– ACAGATAGACGTAGGAGGCATAC – 5’ | This paper | N/A |
| NDUFS3 qPCR  5’– TGGCAGCACGTAAGAAGGG – 3’  3’– CTTGGGTAAGATTTCAGCCACAT – 5’ | This paper | N/A |
| NDUFB2 qPCR  5’– CCCCGGTACAGGGAGTTTC – 3’  3’– GCCAAAATCGCCAAAGAATCCA – 5’ | This paper | N/A |
| SDHA qPCR  5’– GGAACACTCCAAAAACAGACCT – 3’  3’– CCACCACTGGGTATTGAGTAGAA – 5’ | This paper | N/A |
| UQCRC2 qPCR  5’– AAAGTTGCCCCGAAGGTTAAA – 3’  3’– GAGCATAGTTTTCCAGAGAAGCA – 5’ | This paper | N/A |
| COX10 qPCR  5’– AGAAGAGCTATACAGGGATTGCC – 3’  3’– CTGTGTGACATACATGCGCTT – 5’ | This paper | N/A |
| ATP5D qPCR  5’– CCACACTACAGGTCCTACGG – 3’  3’– CACAGAGGAGTCGGCATTCA – 5’ | This paper | N/A |
| TLR9 qPCR  5’– ACAACTCTGACTTCGTCCACC – 3’  3’– TCTGGGCTCAATGGTCATGTG – 5’ | This paper | N/A |
| MYD88 qPCR  5’– AGGACAAACGCCGGAACTTTT – 3’  3’– GCCGATAGTCTGTCTGTTCTAGT – 5’ | This paper | N/A |
| IRF7 qPCR  5’– GCGTACCCTGGAAGCATTTC – 3’  3’– GCACAGCGGAAGTTGGTCT – 5’ | This paper | N/A |
| STAT1 qPCR  5’– TCACAGTGGTTCGAGCTTCAG – 3’  3’– CGAGACATCATAGGCAGCGTG – 5’ | This paper | N/A |
| STAT2 qPCR 5’– GTTACACCAGGTCTACTCACAGA – 3’  3’– TGGTCTTCAATCCAGGTAGCC – 5’ | This paper | N/A |
| CXCL10 qPCR  5’– CCAAGTGCTGCCGTCATTTTC – 3’  3’– GGCTCGCAGGGATGATTTCAA – 5’ | This paper | N/A |
| Primers continued |  |  |
| ISG15 qPCR  5’– GGTGTCCGTGACTAACTCCAT – 3’  3’– CTGTACCACTAGCATCACTGTG – 5’ | This paper | N/A |
| IFIT1 qPCR  5’– GCCTATCGCCAAGATTTAGATGA – 3’  3’– TTCTGGATTTAACCGGACAGC – 5’ | This paper | N/A |
| IFIT3 qPCR  5’– CCTACATAAAGCACCTAGATGGC – 3’  3’– ATGTGATAGTAGATCCAGGCGT – 5’ | This paper | N/A |
| IFI44 qPCR  5’– ATGCTCCAACTGACTGCTCG – 3’  3’– ACAGCAATGCCTCTTGTCTTT – 5’ | This paper | N/A |
| POLG qPCR  5’– GAGCCTGCCTTACTTGGAGG – 3’  3’– GGCTGCACCAGGAATACCAG – 5’ | This paper | N/A |
| TFAM qPCR  5’– GCTCTACACGCCCCTGGTTTCTGG –3’  3’– TCGCTGTAGTGCCTGCTGCTCCTG – 5’ | This paper | N/A |
| TFB2M qCPR  5’– TATAGAGCCGTTGCCTGATTCT – 3’  3’– GCCGCTTTCTTACATGCTATGTG – 5’ | This paper | N/A |
| POLRMT qPCR  5’– AGAAGGCTCCAGTAATGTCCA – 3’  3’– CCTGCATCAGTATGCTCACAA – 5’ | This paper | N/A |
| ENDOG qCPR  5’– TTCCGCGAGGATGACTCTGT – 3’  3’– CACCTGAGGCGCTACGTTG – 5’ | This paper | N/A |
| SSBP1 qPCR  5’– TTCAGTTACTTGGACGAGTAGGT – 3’  3’– CGCCACATCTCATTTGTTGCTA – 5’ | This paper | N/A |
| mtDNA qPCR  5’– CCTATCACCCTTGCCATCAT – 3’  3’– GAGGCTGTTGCTTGTGTGAC – 5’ | This paper | N/A |
| nDNA qPCR  5’– ATGGAAAGCCTGCCATCATG – 3’  3’– TCCTTGTTGTTCAGCATCAC – 5’ | This paper | N/A |
| TNFα qPCR  5’– CAGGCGGTGCCTATGTCTC – 3’  3’– CGATCACCCCGAAGTTCAGTAG – 5’ | This paper | N/A |
| PPARG qPCR  5’– GGAAGACCACTCGCATTCCTT – 3’  3’– GTAATCAGCAACCATTGGGTCA – 5’ | This paper | N/A |
| ANGPTL4 qPCR  5’– GCATCCTGGGACGAGATGAAC – 3’  3’– CCCTGACAAGCGTTACCACA – 5’ | This paper | N/A |
| PDK1 qPCR  5’–GGCGGCTTTGTGATTTGTAT –3’  3’– ACCTAGATCGGGGGATAAAC – 5’ | This paper | N/A |
| VEGFA qPCR  5’– CACCATGCCAAGTGGTCC – 3’  3’– TCTCAATCGGACGGCAGTA– 5’ | This paper | N/A |
| PHD3 qPCR  5’ – GGCCGCTGTACTGTAT – 3’  3’ – TTCTGCCCTTCAGCAT – 5’ | This paper | N/A |
